# Leukemic fitness predicts treatment-free remission and guides adaptive therapy in chronic myeloid leukemia

**DOI:** 10.64898/2026.10.26.762052

**Authors:** Artur C. Fassoni, Nataniel W. Silva, Ingmar Glauche

## Abstract

Chronic myeloid leukemia is controlled by tyrosine kinase inhibitors, yet only about half of well-responding patients who stop therapy remain in long-term remission, and currently it cannot be predicted who will relapse. Mechanistic models reproduce both outcomes but rely on many parameters and longitudinal immune measurements, limiting clinical use. Here we derive a minimal two-equation model in which the immune compartment is eliminated by a quasi-steady-state argument and retained implicitly as a tumor load-dependent kill rate. This reduction lowers the parameter count from fourteen to eight, requires no direct immune measurements, and preserves the two key features of immune-leukemia interaction: a switch-like per-capita kill rate and a bell-shaped immune window peaking at deep remission. A complete analytical treatment shows that a patient’s long-term outcome depends on a single quantity, the leukemic fitness, defined as the ratio of leukemic proliferation to maximal immune-mediated killing. Values above one render the cure steady state unstable, whereas values below one make it stable and therefore attainable; the rate-limiting step is depletion of the quiescent stem cell reservoir below a separatrix, reproducing the clinical link between treatment duration and remission level with durable remission after cessation. Fitted to 75 patients of the DESTINY trial, the minimal model matches the fit quality of its more detailed predecessors. Leukemic fitness estimated from time courses up to treatment cessation, including the dose-reduction period, is associated with outcome, with discrimination improving further when six months of post-cessation data are added. Guided by this single parameter, we design two personalized strategies: extended full-dose therapy for low-fitness patients who have not yet crossed the separatrix, and maintenance dosing for high-fitness patients. Leukemic fitness is thus a clinical predictor that can be estimated from routine monitoring data, and a minimally parametrized model is enough to turn patient trajectories into personalized treatment decisions.

**Author Summary:** Most people living with chronic myeloid leukemia today take a targeted drug that keeps the disease under control, and many who respond well would like to stop treatment. Clinical trials show that about half of them stay healthy without the drug, while the other half relapse and must resume therapy. Today there is no reliable way to tell in advance who belongs to which group. To address this, we built a deliberately simple mathematical model that uses only the routine blood tests already collected during monitoring. From these data, the model weighs how aggressive the leukemia is against how well the immune system keeps it in check and combines the two into a single number we call the leukemic fitness. Using data from 75 patients in a dose-reduction trial, we show that this number separates patients who are likely to stay in remission from those likely to relapse. It also points to personalized options: some patients predicted to relapse could still reach remission if treated a little longer, while for others we can estimate a lower, tailored maintenance dose. In this way, the leukemic fitness could help doctors make better treatment decisions and improve the patient’s quality of life.

## Introduction

Chronic myeloid leukemia (CML) was a fatal malignancy until targeted therapies turned it into a manageable condition [1]. The disease is driven by the Philadelphia chromosome, a reciprocal translocation between chromosomes 9 and 22 that fuses the BCR and ABL1 genes into the BCR::ABL1 oncogene, which encodes a constitutively active tyrosine kinase that sustains the uncontrolled expansion of leukemic cells. Because this fusion gene is unique to the malignant clone, the abundance of its messenger RNA transcript, quantified by real-time quantitative PCR and reported on a standardized international scale, provides a sensitive, near-continuous measure of residual disease and has become the standard for monitoring treatment response [2]. Tyrosine kinase inhibitors (TKIs) targeting BCR::ABL1 achieve deep and durable remissions in most patients, enabling normal life expectancy [1,3]. Optimal therapy response is typically associated with a biphasic decline in BCR::ABL1 levels, in which the first, steeper decline is associated with a reduction in the overall leukemic load, while the slower, secondary decline can be associated with the gradual eradication of more quiescent leukemic stem cells. However, continuous, long-term therapy over many years is associated with side effects, reduced quality of life, and substantial costs [4,5]. Over the past decade, clinical trials have shown that a significant proportion of well-responding patients can safely discontinue TKI therapy and achieve treatment-free remission (TFR) [6–8]. Yet, about half of the patients present with leukemia recurrence within the first year after cessation, highlighting the need for reliable predictors of long-term outcomes [9–11].

Mathematical models have been used to interpret CML treatment dynamics and to explore therapeutic strategies. Earlier models focused on the question of whether the pool of leukemic stem cells (LSC) is targeted by TKIs or not [12–14] and how this process can be intensified by supportive drugs to improve treatment response [15]. Only later, we analytically demonstrated that the biexponential decline of BCR::ABL1 transcripts during TKI therapy can be mechanistically explained by rapid drug-induced killing of proliferating LSCs, followed by a slower decline reflecting rare activation of quiescent LSCs, thereby suggesting reduced TKI doses under maintenance conditions [16]. Complementary approaches incorporated immune effector cells, showing that immunological control is essential to explain sustained remission after TKI discontinuation [17]. In a more recent work we introduced mechanistic detail at the cellular level, linking tumor burden to NK cell number and function and reproducing the concept of an “immune window” [18]. Although these and other models clarified parts of the underlying biology [19–29], no current model can prospectively identify, at or before treatment cessation, which patient will maintain remission, and which will relapse. Closing this gap with predictive models calls for parsimonious approaches preserving the essential mechanisms through a coarser description while remaining identifiable from the BCR::ABL1 measurements routinely collected during patient monitoring.

In this work, we ask whether the mechanisms that govern treatment-free remission can be retained in a model simple enough to be identified from the data actually available in the clinic. Concretely: can the immune compartment, costly to measure and hard to identify, be eliminated without discarding its essential effect on remission? Does the resulting low-dimensional, fully analyzable model remain faithful enough to reproduce individual BCR::ABL1 trajectories? And can it be fitted using only the measurements at hand before a cessation decision, so that a single interpretable quantity captures a patient’s prospects? To address these questions, we translate our previous mechanistic models into a minimal, analytically tractable framework, cutting the parameter count from fourteen to eight, in which the immune response enters implicitly, and the long-term outcome is summarized by a single quantity, the leukemic fitness *ϕ*. The remainder of the paper develops this framework and tests how well such a compact description can anticipate each patient’s remission or recurrence and the treatment adaptations that might be required.

## Methods

The suggested modeling approach builds on a series of progressively refined ODE models developed by our group, each designed to capture distinct aspects of CML treatment dynamics and immune-mediated disease control [16,18,26–28]. Starting from a minimal two-compartment framework for TKI-driven LSC depletion, we successively incorporated immune effector cell populations and, ultimately, mechanistic descriptions of cellular tumor-immune interactions. Here, following the steps of that modeling process, we introduce two mathematically grounded reductions that distill its essential features into a minimal, analytically tractable framework.

### A simple model for CML treatment with TKIs

In [16], we introduced a minimal ODE model to describe the effect of TKIs on LSCs. This formulation was inspired by earlier and more detailed hierarchical models of hematopoiesis [12,13,22] but reduced to the smallest set of parameters. At this stage, no immune effects were considered.

The model describes two LSC populations: quiescent (X) and actively proliferating (Y) cells. The dynamics are governed by the following linear system:

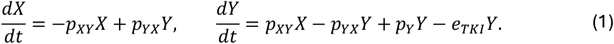

where *p*_*XY*_ and *p*_*YX*_ are the transition rates between quiescent and active states, *p*_*Y*_ is the net proliferation rate of active LSCs, and *e*_*TKI*_ is the TKI-induced kill rate. The model approximates the BCR::ABL1 ratio as *L*_*MOD*_ (*t*) = 100% × *Y*(*t*)/*T*_*Y*_, where *T*_*Y*_ = 10^6^ represents an upper limit for the number of LSC. A linear growth term (*p*_*Y*_*Y*) instead of logistic growth was used for analytical tractability, as the LSC population is consistently decreasing during effective treatment. Using a quasi-steady-state assumption for the initial condition of quiescent LSCs (*X*_0_ = (*p*_*YX*_/*p*_*XY*_)*Y*_0_) and connecting the initial number of active LSCs with the BCR::ABL1 ratio at diagnosis (*Y*_0_ = (*L*_0_/100)*T*_*Y*_), the system reduces to four effective parameters, (*L*_0_, *p*_*XY*_, *p*_*YX*_, *q*).

Clinical BCR::ABL1 time courses of patients responding well under TKI are well described by a bi-exponential decline, obtained phenomenologically by bi-linear regression on a log scale as

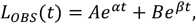

A key feature of the mathematical model (1) is that its analytical solution is also a bi-exponential function:

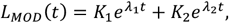

where *λ*_1_, *λ*_2_ are eigenvalues of the system (1),

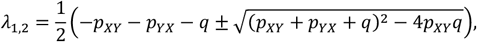

*q* = *e*_*TKI*_ − *p_Y_* is the net balance of TKI-mediated killing and proliferation, and

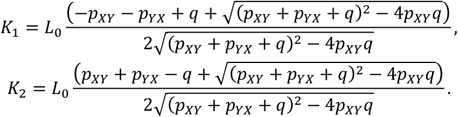

This structure perfectly matches the phenomenological, bi-exponential decline observed in patient data. Thus, by equating the mechanistic model (*L*_*MOD*_) with the phenomenological fit (*L*_*OBS*_), we established an invertible analytical mapping between the two sets of parameters:

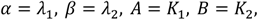

and conversely,

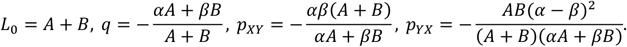

This correspondence allowed for direct estimation of mechanistic parameters from patient-specific bi-exponential fits. Analysis of patient cohorts from the IRIS [3] and CML-IV trials [30] revealed a distinct separation of timescales: the net TKI effect was the fastest (*q* ~ 1 month^−1^), followed by LSC activation (*p*_*XY*_ ~ 0.05 month^−1^) and finally LSC deactivation (*p*_*XY*_ ~ 0.001 month^−1^).

This led to the simplified approximations for the observed slopes:

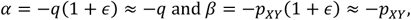

where *ϵ* = *p*_*YX*_/*q* ≈ 0.001. This provided a mechanistic explanation for the bi-exponential decline: an initial, rapid depletion of proliferating LSCs dominated by the net TKI effect (*q*) is followed by a slower decline, which is limited by the rate of quiescent LSC activation (*p*_*XY*_). From this analysis, dose-reduction strategies could be quantitatively assessed. Retrospective fitting suggested that 50% dose reduction would be safe for ~87% of patients achieving MR3, later confirmed by the DESTINY trial [31]. However, this model could not disentangle the individual contributions of the TKI kill rate (*e*_*TKI*_) and the LSC proliferation rate (*p*_*Y*_) from the net effect (*q*).

In summary, this 2-ODE, 4-parameter minimal model captures TKI-driven dynamics: active LSCs are rapidly eradicated, while residual quiescent LSCs persist and fuel slower long-term decline through activation.

### A general model for immunological control of residual disease

While capturing the biphasic decline of BCR::ABL1 transcripts, the minimal TKI model (1) cannot account for treatment-free remission: in the absence of TKI-induced killing, any residual LSC population will inevitably re-expand. Sustained remission after cessation therefore requires an additional suppressive force, which clinical and experimental evidence attributes to immune effector cells [32,33]. To incorporate this, we extended the model by introducing an explicit immune effector cell population, *Z* [26] (Fig 1A):

**Fig 1.**
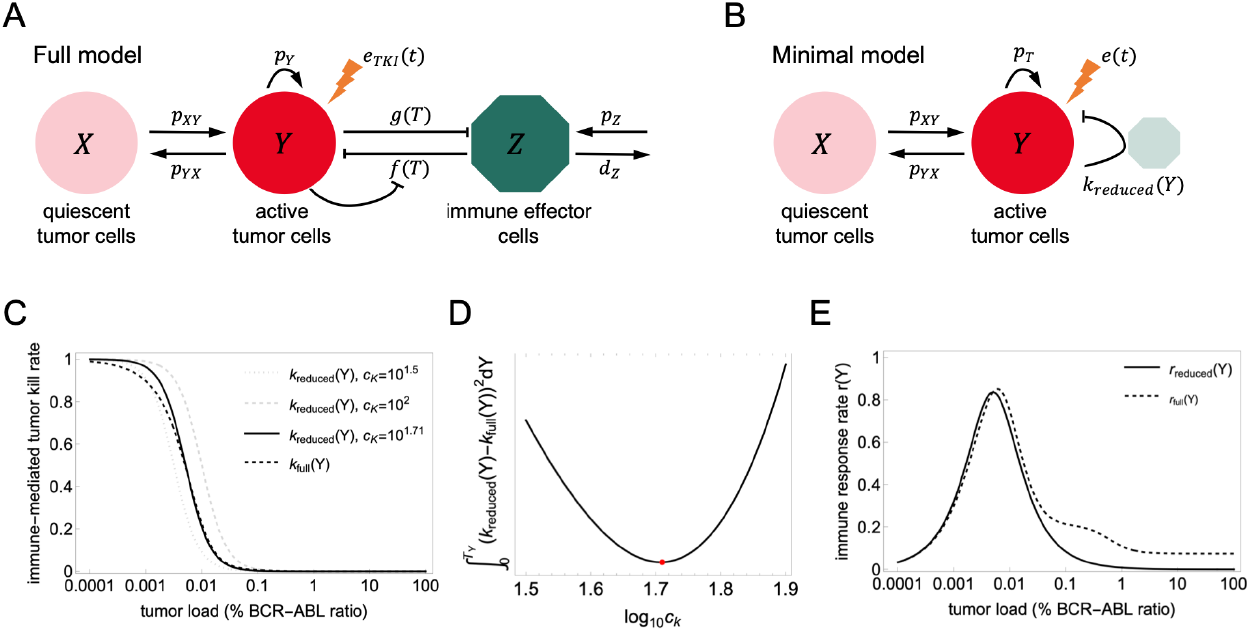
A minimal model for implicit tumor–immune interactions in CML. **(A)** Sketch of the detailed tumor–immune model (Methods), comprising active leukemic stem cells (*Y*), quiescent niche-bound leukemic stem cells (*X*), and immune effector cells (*Z*). Active LSCs proliferate, transition to and from the quiescent compartment, are eliminated by TKI treatment, and interact with immune effector cells. Immune cells are produced at a constant rate, die naturally, and their number and function are modulated by tumor load. **(B)** Sketch of the minimal model, in which the immune compartment is eliminated via a quasi-steady-state approximation. All immune effects are implicitly encoded in the per-capita kill rate *k*_*reduced*_(*Y*), reducing the system to two ODEs for *Y* and *X*. The faded immune cell icon reflects the implicit, rather than explicit, representation of immune dynamics. **(C)** Comparison of the mechanistically derived per-capita immune kill rate *k*_*full*_(*Y*) (dashed line) with *k*_*reduced*_(*Y*) for three values of *C*_*K*_. Both functions exhibit a switch-like profile, transitioning from near-zero at high tumor loads, where immune suppression dominates, to a plateau at low tumor loads where immune activity is restored. **(D)** Integrated squared error between *k*_*reduced*_ (*Y*) and *k_full_*(*Y*) as a function of log_10_ *C_K_*, minimized (red dot) at *C*_*K*_ = 10^1.71^. This value is adopted as a fixed population-level parameter throughout the analysis. **(E)** Comparison of the systemic immune response rates *r*_*reduced*_ = *k*_*reduced*_ (*Y*)*Y* with *C*_*K*_ = 10^1.71^ (solid line) and *r*_*full*_(*Y*) (dashed line). Both functions display a bell-shaped immune window, with a peak near MR4 (0.01% BCR::ABL1), reflecting maximal antitumor immune activity at intermediate-to-low tumor loads. The close agreement between the two curves, particularly in the low tumor load region relevant for TFR dynamics, substantiates the proposed reduction.

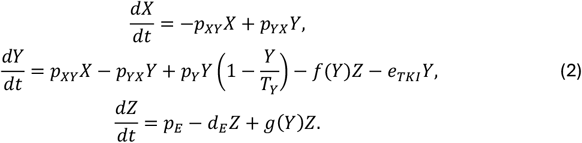

Here, *p*_*E*_ and *d*_*E*_ denote the basal production and natural death rates of immune effector cells, respectively. The term *f*(*Y*)*Z* captures immune-mediated killing of LSCs, while *g*(*Y*)*Z* models the modulation of immune cell numbers by tumor load (either recruitment at intermediate or suppression at high burdens). To avoid unbounded solutions in the mathematical analysis, a logistic growth term is used for active LSCs. The functional forms *f*(*Y*) and *g*(*Y*) are not specified a priori; instead, we systematically explored 4 choices for *f*(*Y*) and 5 for *g*(*Y*), yielding 20 candidate submodels [26]. Although many submodels fitted patient data comparably well, qualitative analysis of their phase portraits favored formulations with biologically plausible equilibrium structures.

One such submodel was applied to patient data after TKI cessation [28], reproducing three clinically observed outcomes: (i) deep remission with undetectable disease, (ii) stable low-level remission with persistent but controlled disease, and (iii) molecular relapse. In that model, we adopted a bell-shaped functional response for the immune killing term,

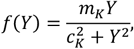

known as the immune window [17]. This form reflects the non-monotonic nature of anti-tumor immunity: killing efficiency increases with tumor load at low burdens, peaks at *Y* = *C*_*K*_, and declines at high burdens due to tumor-induced immune suppression. This bell-shaped form was favored by phase portrait analysis and later derived mechanistically from first principles of cellular interactions, as detailed in the following subsection.

Despite its biological appeal, fitting this model to pre-cessation time courses revealed an inherent limitation: multiple parameter sets could reproduce the same BCR::ABL1 trajectory yet predict divergent outcomes upon treatment withdrawal [27]. This non-uniqueness reflects the structural challenge of estimating immune parameters from on-treatment data alone, and motivates the more mechanistic approach described in the following subsection.

### A detailed model for tumor-immune cellular interactions

While the above, general model established that immune-mediated control is essential for TFR, its functional forms *f*(*Y*) and *g*(*Y*) remained phenomenological. To place these interactions on a mechanistic basis, we developed a refined model in which the immune killing term emerges from first principles of cellular dynamics [18].

Specifically, we modeled the sequential stages of an immune effector cell’s anti-tumor “incursion”: (i) a search phase, during which the cell scans the tissue for a tumor target; (ii) a handling phase, encompassing target recognition, binding, and killing; and (iii) a recharge phase, during which the cell recovers full cytotoxic functionality before engaging again. Combining this kinetic description with a tumor load-dependent suppression of immune cell functionality yields a composite per-capita killing rate,

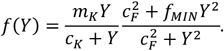

The first factor is a Holling Type II saturating term and captures the density-dependent searching-handling-recharging process where killing efficiency increases with tumor load due to more frequent encounters but saturates as immune cells become occupied with handling. The second factor accounts for tumor-induced suppression of immune cell functionality: at low tumor burdens, immune cells operate near full capacity, while increasing tumor load progressively reduces the fraction of functional cells, down to a minimum *f*_*MIN*_. Together, these two mechanisms produce a composite per-capita killing rate that is bell-shaped: it peaks at intermediate tumor loads where encounters are frequent and immune function is still preserved and declines at both extremes.

In the same model, the modulation of immune cell numbers was described by a suppressive interaction consistent with our earlier modeling [26],

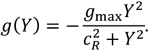

This formulation was fitted to patient data containing paired BCR::ABL1 time courses and longitudinal NK cell number and function measurements from eight patients [10], capturing the joint evolution of tumor and immune compartments during TKI treatment.

### Timescale separations in tumor-immune interactions

An important structural property of the previous model is a clear separation of timescales: immune cell dynamics operate on a faster scale than LSC dynamics. This justifies applying a quasi-steady-state approximation (QSSA) to the immune compartment, setting *dZ*/*dt* = 0 in (2) to obtain the tumor load-dependent equilibrium:

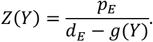

Substituting *Z*(*Y*) into the LSC equations yields a reduced two-dimensional system:

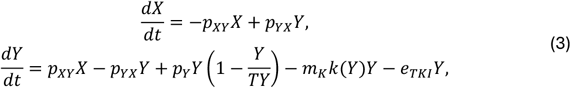

in which all immune effects are encoded in the composite per-capita kill rate

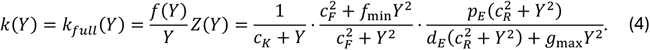

This function exhibits a switch-like profile: it is nearly zero at high tumor loads, where immune suppression dominates; rises sharply as tumor burden falls below a critical threshold; and plateaus at low tumor levels, where immune cells operate at maximal efficiency (Fig 1C). The systemic rate *r*_*full*_(*Y*) = *Yk*_*full*_(*Y*),

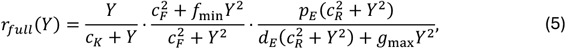

then recovers the bell-shaped immune window as a collective property of the model (Fig 1E).

This timescale separation was verified numerically across a range of parameter values [18], confirming that the slow manifold reduction, eqs. (3-4), provides an accurate description of the model immune cell dynamics independent of the specific parameter values.

Despite this mechanistic richness, the identifiability limitation remained. The model fitting was implemented with a stepwise strategy: first estimating LSC parameters (*L*_0_, *p*_*XY*_, *p*_*XY*_, *q*) from the biphasic decline, exactly as the fitting of model (1), then estimating immune parameters (*c*_*K*_,*c*_*F*_, *f*_*min*_, *p*_*E*_, *d*_*E*_, *g*_*max*_, *c*_*R*_, *Z*(0)) from NK cell time courses, and finally the remaining parameters (*p*_*Y*_, *m*_*K*_) from the complete trajectory. Yet even with this scale-separation-based approach, these last two parameters, the intrinsic LSC proliferation rate *p*_*Y*_ and the maximum immune kill rate *m*_*K*_, resist unique estimation. Their opposing effects are effectively masked by the dominant action of TKI during treatment and can only be disentangled by observing the system’s response to perturbations, such as dose reduction or treatment cessation. This persistent non-identifiability, even in a mechanistically detailed model, directly motivates the reduced formulation introduced below.

## Results

### A minimal model for implicit tumor-immune interactions

In our previous line of ODE models for CML (Fig 1A, Methods), we progressively increased physiological detail, starting with a TKI-only model (equation (1)), to the general tumor– immune interaction model (equation (2)), and finally to the detailed cellular-level formulation [18], and its QSSA-reduced form (equation (3)). Both the general and detailed models demonstrated that immune-mediated control is essential to explain treatment-free remission but also revealed a structural limitation in parameter identifiability: the balance between leukemic cell growth (*p*_*Y*_) and immune-mediated killing (*m*_*K*_) remains unresolvable from on-treatment data alone. Furthermore, we showed that the behavior of the detailed model was equivalent to the formulation obtained after time-scale separation when immune effector cells were assumed to be in a quasi-steady state. However, even this amended model formulation still retained 14 parameters. Motivated by these findings, we now introduce a minimally parametrized model that implicitly incorporates immune effects (Fig 1B) and retains only the essential features of tumor–immune interactions while substantially reducing parameter complexity.

The starting point for this functional reduction is the QSSA-based two-dimensional system derived for the detailed model in Methods (equation (3)), in which immune effector cells no longer appear as an explicit compartment but are encoded in the composite per-capita kill rate *k*_*full*_(*Y*) (equation (4)). This function, derived from NK cell data and cellular-level dynamics, has seven parameters and a complex analytical form. However, its essential qualitative feature is a switch-like profile: *k*_*full*_(*Y*) is nearly zero at high tumor loads, where immune suppression dominates, and rises sharply to a plateau as tumor burden falls below a critical threshold (Fig 1C, dashed line). This emergent shape motivates a phenomenological substitution: rather than carrying the full mechanistic complexity of *k*_*full*_(*Y*), we ask whether a much simpler one-parameter function can reproduce the same qualitative behavior.

We propose the following reduced per-capita immune kill rate (Fig 1C):

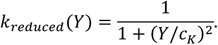

This expression is switch-like by construction: it approaches 1 at low tumor loads (immune activity fully restored), equals 0.5 at *Y* = *C*_*K*_, and vanishes at high tumor loads (immune suppression). The single parameter *C*_*K*_ defines the tumor load at which immune efficacy is halved; we therefore refer to *C*_*K*_ as the half-efficacy tumor load. To identify the value of *C*_*K*_ that best approximates the mechanistically derived *k*_*full*_(*Y*), we minimize the integrated squared error between the two functions (Fig 1D), obtaining an optimal value of *C*_*K*_ = 10^1.71^. This value will be used as a fixed, population-level constant throughout the analysis; the effect of varying *C*_*K*_ over several orders of magnitude on the kill rate and on the bifurcation structure is shown in S1 Appendix Fig B. Because this mechanistic kill rate is not measured directly but results from a complex, multistep derivation based on the full model and its calibration to patient data, the half-efficacy tumor load should be read as a compact summary of that derivation. The exponent 2 in the denominator is a modeling choice; it controls how abruptly immune efficacy is lost as tumor load rises, with larger exponents producing a steeper transition around *C*_*K*_and, in the limit, a step function. We adopt 2 as the smallest and simplest exponent that reproduces the switch-like profile of the mechanistically derived kill rate while keeping the equilibrium condition a cubic polynomial in *Y*, which makes the complete bifurcation analysis (presented below) analytically tractable.

Substituting *k*(*Y*) = *k*_*reduced*_(*Y*) instead of *k*_*full*_(*Y*) in the full model (equation (3)) reduces the parameter count by 6 and leads to the minimal two-dimensional system (Fig 1B)

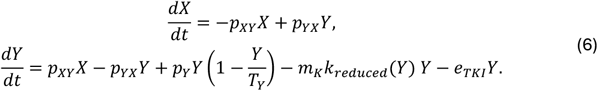

The term *m*_*K*_*k*_*reduced*_(*Y*)*Y* encapsulates all underlying immunological processes, from cellular-level interactions to systemic suppression, into a single functional form, with *m*_*K*_ representing the maximum immune-mediated kill rate. The corresponding systemic immune response rate, *r*_*reduced*_ (*Y*) = *Yk*_*reduced*_ (*Y*), recovers the bell-shaped immune window (Fig 1E, solid line), with its maximum occurring near MR4 (0.01% BCR::ABL1), and closely matches the shape of *r*_*full*_(*Y*) derived from the full model (Fig 1E, dashed line). Although the two curves deviate between 0.1% and 1% BCR::ABL1 with the full model producing a slower descent, the per-capita kill rates align closely over the same range (Fig 1C). Furthermore, this deviation occurs above the tumor loads at which the tumor-immune balance and the fate of treatment-free remission are commonly determined. Therefore, it has minimal to no impact on subsequent results.

In summary, this minimal model distills the immune-mediated kill process into a single parameter *C*_*K*_, reducing the total parameter count from 14 to 8. It preserves the two essential qualitative features of the detailed model: the switch-like per-capita kill rate and the bell-shaped immune window. In the following sections, we show that this reduction retains the same fitting ability as the detailed model, without explicitly requiring longitudinal immune cell measurements, while still keeping the identifiability challenge of disentangling *p*_*Y*_ from *m*_*K*_.

### Fit quality and parameter identifiability with complete time courses

Having established the minimal model as a mathematically consistent simplification of the detailed tumor–immune framework, we next assessed its practical performance in two complementary ways. First, we compared the quality of model fits to patient time courses against those of the full model; and second, we evaluated the model’s ability to predict treatment outcomes.

We applied the minimal model to BCR::ABL1 time courses of 75 patients from the DESTINY trial [31], the same cohort used for model fitting in [18] and in earlier work of our group [27], where the patients were selected on the basis of follow-up duration and a sufficient number of available measurements; no further exclusions were applied here. Each patient underwent initial full-dose TKI therapy (median 6.9 years, interquartile range 4.8–10.2 years, in the full DESTINY cohort) [31]. Eligibility for dose reduction required at least three years of TKI therapy and all BCR::ABL1 measurements over the preceding 12 months at or below 0.1% (MR3); patients whose measurements were additionally all at or below 0.01% (MR4) formed the stable-MR4 group, the remainder the MR3-but-not-MR4 group. Upon trial entry TKI dose was reduced to 50% of the recommended full dose for 12 months before subsequent treatment cessation [31].

For the comparison that follows, we briefly review the fitting strategy for the full model [18], which was kept identical for the minimal model, differing only in how the immune kill rate *k*(*Y*) is parametrized. In both models the carrying capacity is fixed at *T*_*Y*_ = 10^6^ without loss of generality, and the four LSC parameters (*L*_0_, *p*_*XY*_, *p*_*YX*_, *q*), where *q* = *e*_*TKI*_ − *p*_*Y*_ is the net TKI effect, are obtained first: since the immune response is negligible relative to the TKI effect before MR4, the full and immune-free models coincide there, so the biexponential *L*_*OBS*_(*t*) = *Ae*^*αt*^ + *Be*^*βt*^ is fitted to the full-dose data above MR4 and the four parameters recovered from (*A, B, α, β*) through their analytical correspondence (see Methods). The two models then differ only in the fixed parameters of the immune kill rate: the full model carries eight immune parameters (*C*_*K*_, *c*_*F*_, *f*_*min*_, *p*_*E*_, *d*_*E*_, *g*_*max*_, *c*_*R*_, *Z*_0_), which are not identifiable from the DESTINY data (no longitudinal NK-cell measurements) and are therefore fixed at the values estimated from the median NK-cell time courses of the eight patients in [10]; the minimal model collapses these into the single half-efficacy tumor load *C*_*K*_, fixed at the population value 10^1.71^ cells so that its reduced kill rate *k*_*reduced*_ matches the full-model kill rate *k*_*full*_ (Fig 1C, Methods). With the kill rate thus fixed, both models are left with exactly the same two free parameters, the tumor growth rate *p*_*Y*_ and the maximum immune kill rate *m*_*K*_, estimated from the complete patient trajectory (full dose, then half dose, then off-treatment) by the same Monte Carlo procedure used in [18]: 10^4^ pairs are drawn uniformly from [0, 3] for *p*_*Y*_ and [0, 6] for *m*_*K*_ (both in month^−1^), ranked by the residual sum of squares over all of a patient’s measurements, and the 100 best fits retained. Using identical sampling ranges and sample size for both models allows a direct comparison of fit quality and predictive performance while preserving the core identifiability challenge, namely disentangling the antagonistic effects of *p*_*Y*_ and *m*_*K*_, and, for the minimal model, entirely bypassing the need for longitudinal immune-cell measurements.

Comparing fit quality between the two models reveals that the minimal model performs at least as well as the full model across the patient cohort. The distribution of per-patient residuals is comparable, with the minimal model showing a slightly lower median and a more compact spread (Fig 2A, left). A patient-level scatter plot confirms that for most patients, residuals from the minimal model fall below those of the full model, lying beneath the diagonal (Fig 2A, right). This indicates that the functional simplification of the immune kill rate does not compromise the descriptive accuracy of the model, and in several cases marginally improves it. For interpreting this result, we point out that the additional seven immune parameters of the full model are not free but fixed at values transferred from a different cohort (see Methods), while the single immune parameter of the minimal model is fixed at a value matching the resulting kill rate of the full model (Fig 1C,D). Ultimately, both models have the same free parameters, while differences in fit quality between them emerge from the *shape* of their kill rates.

**Fig 2.**
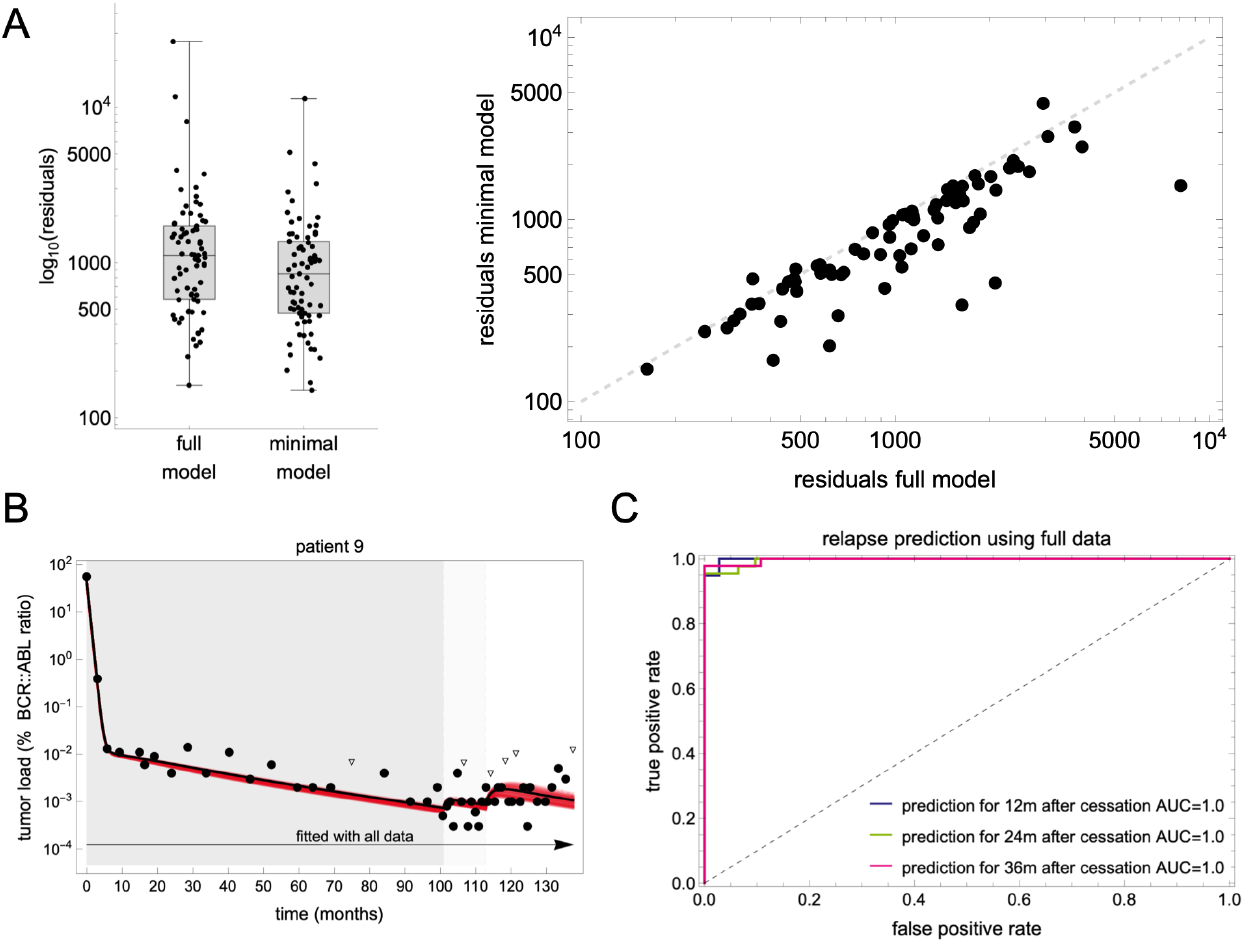
Model fitting quality and recurrence prediction using complete time courses. **(A)** Comparison of fitting residuals between the full model [18] and the minimal model across the 75 DESTINY patients. Left: distribution of per-patient residuals (log_10_ scale) shown as boxplots with individual data points; the minimal model shows a slightly lower median and more compact spread. Right: scatter plot of per-patient residuals from the minimal model versus the full model; points below the diagonal (dashed line) indicate better fits by the minimal model, which accounts for most patients. **(B)** Model fitting and recurrence prediction for a representative patient (patient 9) using the complete time course. Left: the 100 best-fit trajectories (red) and best overall fit (black) are shown alongside BCR::ABL1 measurements (filled circles: detected; open triangles: undetectable). The dark and light gray shaded regions indicate the full and half dose periods, respectively. Fits are tightly concentrated, reflecting good parameter identifiability when all data are available. **(C)** Receiver operating characteristic (ROC) curves for predicted molecular recurrence at 12, 24, and 36 months after cessation, based on the fraction of the 100 best fits predicting tumor load above MR3. The AUC = 1.0 across all time horizons indicates that the retained fits agree with the *reference outcome* on the direction of the long-term outcome for every patient; since the *reference outcome* is itself the best member of this ensemble, the value reflects internal coherence rather than validation against an independent endpoint.

To compare model fits with patients’ true outcomes, we distinguish two outcome definitions. The first is the *clinical outcome*, which classifies a patient as having disease recurrence when the *measured* BCR::ABL1 ratio exceeds 0.1% (loss of MR3) during follow-up after cessation. The second, named the *reference outcome*, corresponds to the simulated BCR::ABL1 ratio exceeding 0.1%, computed using the best-fit parameter values obtained from the complete time course. Because the simulated *reference outcome* reproduces the measured *clinical outcome* for all 75 patients, the two definitions coincide. The *clinical outcome*, however, cannot be evaluated at a common horizon because follow-up times vary widely across the cohort, and for many patients the last measurement falls only a few months after cessation, in several cases less than six. Consequently, predictions from different truncation windows cannot be compared at fixed horizons such as 36 months after treatment cessation. We therefore adopt the *reference outcome* as a surrogate because, being a model solution, it can be extrapolated beyond the last measurement to a horizon shared by all patients.

When fitted to the *complete time course* of each patient (up to the time of the last available measurement, denoted by *t*_*F*_), the 100 best parameter pairs produce highly consistent trajectories, with the ensemble of fits converging tightly around the observed data (Fig 2B). Reflecting the non-uniqueness of *p*_*Y*_ and *m*_*K*_, these fits span a range of post-cessation predictions; however, when used to classify patients by predicted molecular recurrence at different time points after cessation (*t_F_* + 12, *t_F_* + 24, *t_F_* + 36), the model achieves perfect discrimination, with AUC = 1.0 across all time horizons (Fig 2C). Because the score being thresholded (the fraction of the ensemble predicting recurrence) and the label being predicted (the outcome of the ensemble’s own best member) both derive from the same fit, this value assesses internal coherence and establishes only that the ensemble is not degenerate at the last measurement. The residual spread in the proliferation and immune kill rates is small enough that every retained fit agrees with the optimum on the direction of the long-term outcome, for each patient. The retained ensemble also makes the geometry of this non-uniqueness explicit: within each patient, the 100 best parameter pairs do not scatter without a preferred direction but rather lie along a narrow ridge where a higher proliferation rate is balanced by a higher immune kill rate (S1 Appendix Fig A). The within-patient rank correlation between the two rates is positive in every patient, with median *ρ* = 0.84 (interquartile range 0.66–0.93), and significant at the 5% level in 74 of 75 patients.

We conclude that the minimal model matches the descriptive accuracy of the full model while depending on fewer parameters and no longitudinal immune-cell measurements. This leads to the practical question addressed next: how much of the trajectory must be observed before a patient’s long-term outcome can be anticipated.

### Predicting treatment-free remission from truncated time courses

Having established that the complete time course carries the information needed to predict the outcome, we next asked how much of it is actually required. We therefore refitted the minimal model to progressively truncated trajectories, removing data from the end backwards, which mimics the clinical scenario in which a prediction must be made before the outcome is observed. Following the approach of [18], we considered three truncation windows: data up to six months after cessation (*t_C_* + 6), data up to the time of treatment cessation (*t_C_*), and data up to the start time of dose reduction (*t_R_*).

Truncating the trajectory at six months after cessation (up to *t_C_* + 6) retains almost all the predictive information, with AUC values of 0.94–0.97 across all time horizons (Fig 3A). The early post-cessation dynamics, in which the immune system is directly challenged by residual tumor cells in the absence of TKI, provide the decisive information needed to resolve the tumor-immune balance.

**Fig 3.**
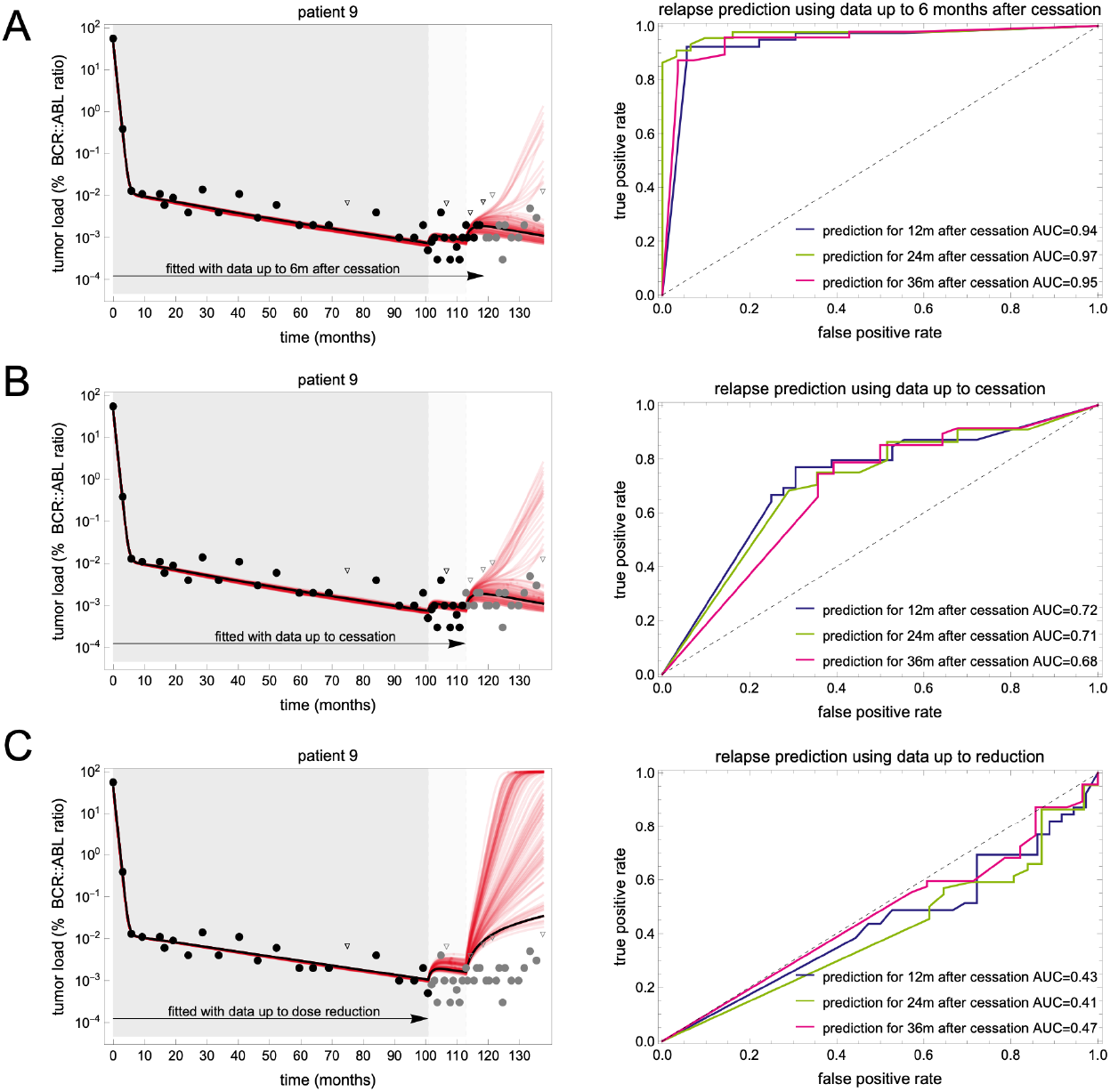
Prediction of treatment-free remission using progressively truncated data. Each row corresponds to a different truncation window, in order of decreasing information, with the left panel showing individual model fits for patient 9 and the right panel showing ROC curves across the 75 DESTINY patients. In each left panel, black circles indicate data used for fitting, triangles indicate undetectable values used for fitting, and gray points indicate data withheld from fitting. The dark and light gray shaded regions indicate the full and half dose periods, respectively. ROC curves show recurrence classification performance at 12 months (blue), 24 months (green), and 36 months (magenta) after cessation. **(A)** Fitting with data up to six months after cessation (*t_C_* + 6*m*). Trajectories converge substantially and recurrence prediction improves markedly, reaching AUC ≈ 0.95. The early post-cessation dynamics, in which the immune system directly confronts residual tumor cells in the absence of TKI, provide the decisive signal needed to resolve the tumor–immune balance and enable reliable individual-level predictions. **(B)** Fitting with data up to treatment cessation (*t_C_*), including the dose reduction period. Trajectories diverge more after cessation than in (A), and recurrence prediction decreases to AUC ≈ 0.70, indicating that the kinetics of tumor response to dose reduction carry partial but incomplete predictive information. **(C)** Fitting with data up to dose reduction (*t_R_*). The 100 best-fit trajectories diverge substantially after cessation, reflecting the fundamental non-identifiability of *p_Y_* and *m_K_* from full-dose data alone. Recurrence prediction is not informative, with AUC below 0.50 across all time horizons.

Truncating further, at the time of treatment cessation (up to *t_C_*) removes that decisive signal, and predictive performance falls to AUC values of approximately 0.68-0.72 (Fig 3B). The kinetics of tumor load response to dose reduction still provide partial information on the *p_Y_*/*m_K_* balance, constraining the parameter space but increasing trajectory divergence after cessation, though substantial uncertainty remains.

Finally, discarding the dose reduction period as well and fitting data under full dose only (up to *t_R_*), the 100 best fits produce highly divergent post-cessation trajectories, reflecting the fundamental non-identifiability of *p_Y_* and *m_K_* from on-treatment data alone (Fig 3C, left). Accordingly, the resulting recurrence predictions are essentially uninformative, with AUC values below 0.50, indistinguishable from random classification (Fig 3C, right). This confirms, in the context of the minimal model, the finding of [18] that BCR::ABL1 dynamics under full-dose therapy carry no useful predictive signal for TFR.

Together, these results show that the minimal model reproduces, across all truncation windows, the predictive pattern reported for the full model in [18] (AUCs equal to 0.53, 0.71, 0.92 for prediction 36 months after cessation using data up to *t_R_, t_F_* and *t_F_* + 6 respectively), with fewer parameters, and requiring neither longitudinal immune cell measurements nor a dedicated second fitting step.

### Insights from stability analysis and bifurcation structures

A key advantage of the minimal model is its analytical tractability, which enables a complete characterization of its equilibrium structure and bifurcation behavior. This analysis helps interpret the origin of the parameter non-identifiability outlined above and shows what treatment is needed to achieve immune-mediated cure.

Setting *dX*/*dt* = *dY*/*dt* = 0, the equilibria satisfy *X* = (*p_YX_*/*p_XY_*)*Y* and

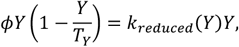

where *ϕ* = *p_Y_*/*m_K_* is the sole bifurcation parameter, the ratio of the leukemic proliferation rate to the maximum immune kill rate, which we term the *leukemic fitness*. One solution is always *Y* = 0, corresponding to the cure equilibrium *E*_0_ = (0,0). As shown in S1 Appendix, *E*_0_ is stable when *ϕ* < 1 (low leukemic fitness) and becomes a saddle when *ϕ* > 1, in which case mathematical cure is unattainable regardless of treatment duration.

The remaining equilibria are roots of the cubic polynomial

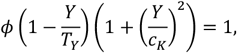

which admits up to three positive roots in general. In the biologically relevant parameter regime (*ϕ* < 1, low leukemic fitness), at most two positive roots exist, *Y_H_* > *Y_L_* > 0, giving rise to a stable, high-load equilibrium *E_H_* (corresponding to disease recurrence) and a low-load equilibrium *E_L_* (a saddle point defining the boundary between the basins of attraction of *E*_0_and *E_H_*). A third root near *Y* = 0 can arise for *ϕ* slightly above 1, but the corresponding equilibrium lies below the threshold of a single cell and is therefore biologically irrelevant. We also prove that no periodic solutions occur and all trajectories converge either to *E*_0_ or *E_H_* (see proofs and details in S1 Appendix, including Fig B therein).

The resulting bifurcation diagram (Fig 4A) summarizes the global dynamics as a function of the leukemic fitness *ϕ*. At very low *ϕ, E*_0_ is the only attractor, implying complete disease control. As *ϕ* increases, a saddle-node bifurcation creates *E_L_* and *E_H_*, establishing bistability: the stable manifold of *E_L_* acts as a separatrix partitioning the phase plane into the basins of attraction of *E*_0_ (cure) and *E_H_* (recurrence). As *ϕ* approaches 1 from below, *Y_L_* decreases, implying that the boundary of the cure basin shrinks toward zero, until a transcritical bifurcation occurs at *ϕ* = 1 after which *E*_0_ becomes unstable and *E_H_* globally attracting.

**Fig 4.**
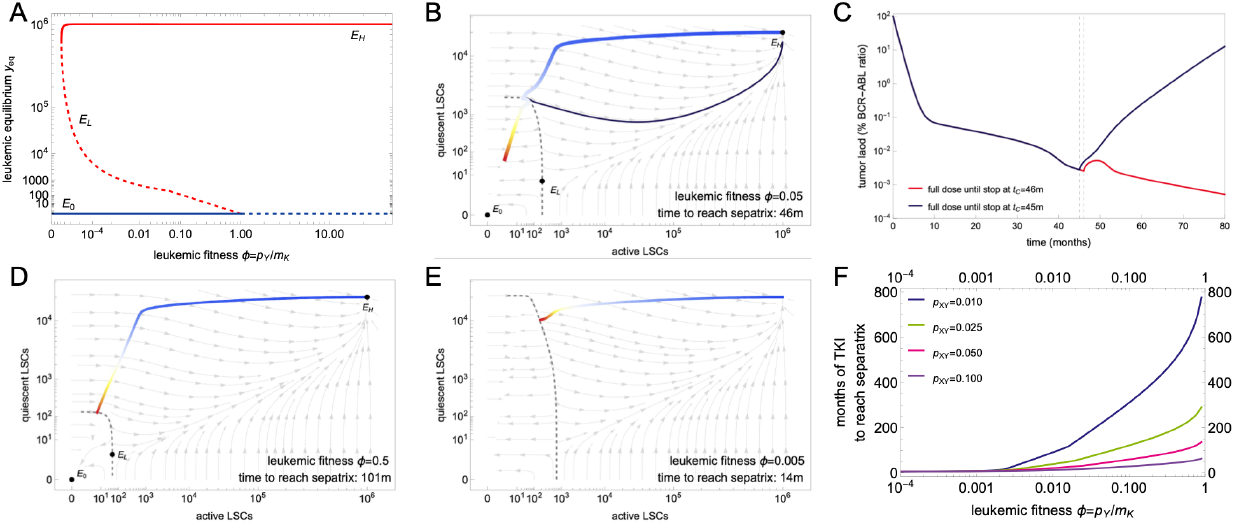
Bifurcation structure and treatment implications of the minimal model. **(A)** Bifurcation diagram showing the steady-state leukemic cell counts as a function of leukemic fitness *ϕ* = *p_Y_*/*m_K_*. Three regimes are identified: for *ϕ* below a saddle-node threshold, only the cure equilibrium *E*_0_ exists and is stable (solid blue line); for *ϕ_SN_* < *ϕ* < 1, bistability arises between the cure equilibrium *E*_0_ (solid blue line) and the high-load disease equilibrium *E_H_* (solid red line), with the saddle point *E_L_* defining the separatrix (dashed red line); for *ϕ* > 1, *E*_0_ loses stability (dashed blue line) and *E_H_* becomes the global attractor (solid red line). **(B, D, E)** Phase portraits of the minimal model for three values of leukemic fitness: *ϕ* = 0.05 **(B)**, *ϕ* = 0.5 **(D)**, and *ϕ* = 0.005 **(E)**. Gray arrows indicate the vector field of the system without treatment (*e_TKI_* = 0); the dashed curve marks the separatrix (stable manifold of *E_L_*). Colored trajectories represent simulated patient dynamics under full-dose TKI treatment, with color encoding time (blue: early treatment; white/yellow: intermediate; red: late treatment/cessation). The time required to cross the separatrix and enter the basin of attraction of cure is 46, 101, and 14 months for panels B, D, and E, respectively. Both axes are shown on a logarithmic scale. **(C)** Corresponding BCR::ABL1 time courses for the phase portrait in **(B)**, illustrating the critical dependence on treatment duration. Cessation at *t_C_* = 46 months (colored solid line), after the trajectory has crossed the separatrix, leads to sustained remission; cessation at *t_C_* = 45 months (dark blue solid line), before crossing, results in molecular recurrence. **(F)** Time required to reach the separatrix under full-dose TKI therapy as a function of leukemic fitness *ϕ*, for four values of the quiescent LSC activation rate *p_XY_*. Treatment duration requirements increase steeply as *ϕ* approaches 1 and decrease with higher activation rates due to faster depletion of the quiescent reservoir. The value *p_XY_* = 0.050 month^−1^ (magenta) is the reference value and was used in panels A-E.

This bifurcation structure has direct implications for understanding treatment dynamics. Fig 4B illustrates the phase portrait and a representative patient trajectory under full-dose TKI therapy, with leukemic fitness *ϕ* = *p_Y_*/*m_K_* = 0.05. The initial steep decline in active LSCs (*Y*, horizontal movement in phase space, blue segment) corresponds to the fast first slope of BCR::ABL1 decline, driven primarily by TKI cytotoxicity. The subsequent diagonal phase reflects the slow release of niche-bound quiescent LSCs (*X*) into the active compartment where they are subject to treatment (white segment), thereby corresponding to the second, shallower slope of CML response kinetics. Whether the trajectory crosses the separatrix, and thus enters the basin of attraction of *E*_0_, depends critically on how long treatment is maintained. For this representative patient, cessation at or before 45 months results in recurrence (dark blue trajectory and Fig 4C), while cessation at 46 months leads to cure. It is the quiescent LSC reservoir that governs this sharp transition: even when active LSC numbers fall near the threshold *Y*_*L*_, the quiescent pool *X* remains elevated and sustains reseeding of the active compartment.

The time required to cross the separatrix depends strongly on leukemic fitness *ϕ* and on the quiescent cell activation rate *p*_*XY*_ (Fig 4D-F). Increasing *ϕ* tenfold (*ϕ* = 0.5) shrinks the cure basin substantially, requiring 101 instead of 46 months of full-dose therapy to reach the separatrix (Fig 4D). Conversely, decreasing *ϕ* tenfold (*ϕ* = 0.005) rapidly enlarges the cure basin; the separatrix becomes nearly vertical, the quiescent reservoir depletes quickly, and only 14 months suffice (Fig 4E). Fig 4F further quantifies this dependence across a range of *ϕ* and *p*_*XY*_ values: for leukemic fitness near *ϕ* ≈ 0.1 (tenfold below the curability threshold) treatment requirements remain substantial, and they even reach hundreds of months once the quiescent activation rate *p*_*XY*_ is lowered from its default value of 0.05 month^−1^ (magenta curve) to 0.025 and 0.01 month^−1^ (green and dark blue curves). In such cases, the required treatment durations are clinically feasible only for very low leukemic fitness. Conversely, higher activation rates *p*_*XY*_ accelerate depletion of the quiescent pool, thereby substantially reducing the time to cross the separatrix for any given *ϕ* (*p*_*XY*_ = 0.1 month^-1^, purple curve).

Together, these results provide a mathematical explanation for the well-established clinical observation that deep and sustained molecular response is a prerequisite for successful treatment-free remission [34]: it is not sufficient to suppress active disease, the quiescent LSC reservoir must be depleted below the separatrix threshold, a process that is inherently slow and patient-specific [15]. The same mechanism explains why total time on TKI and, to a lesser extent, time spent in MR4 are among the strongest clinical predictors of successful cessation [34,35]: in the model, both are proxies for how far the quiescent reservoir has been depleted toward the separatrix.

### Predicting treatment response from estimated leukemic fitness

The above mathematical analysis established that the leukemic fitness *ϕ* = *p*_*Y*_/*m*_*K*_ is the single parameter governing the asymptotic behavior of the system: for *ϕ* < 1, the cure equilibrium *E*_0_ is stable and sustained remission is achievable; for *ϕ* > 1, *E*_0_ is unstable and recurrence is the inevitable long-term outcome regardless of treatment duration. Estimating the parameter pairs (*p*_*Y*_, *m*_*K*_) from patient time courses therefore amounts to estimating the leukemic fitness *ϕ*, and the bifurcation structure immediately translates model fits into outcome predictions with clinical relevance.

We now evaluate the predictive value of fitness-based classification. Technically, we obtain the best fit (*p*_*Y*_, *m*_*K*_) for each patient and each data selection strategy (up to *t*_*R*_, *t*_*C*_, *t*_*C*_ + 6 and *t*_*F*_), compute the leukemic fitness *ϕ* = *p*_*Y*_/*m*_*K*_ and compare it against the *reference outcome* at 36 months after cessation (*t*_*C*_ + 36). Stratifying patients into recurrence and remission groups according to the reference outcome reveals that the distributions of fitness values *ϕ* for these groups separate progressively as more data are incorporated (Fig 5A; two-sided Mann–Whitney U test throughout): the separation is not statistically significant when using data up to *t*_*R*_ (*p* = 0.173), becomes marginal up to *t*_*C*_ (*p* = 0.051), and reaches high significance with data up to *t*_*C*_ + 6 and *t*_*F*_ (*p* = 1.4 × 10^−4^ and *p* = 9.6 × 10^−6^, respectively). The same trend holds when the mean fitness across all 100 best fits is used instead of the single best fit (Fig C in S1 Appendix). We note that for *t*_*F*_ the fitness and the *reference outcome* are both derived from the same best fit, so this window quantifies internal consistency and serves as an upper reference for the truncated windows.

**Fig 5.**
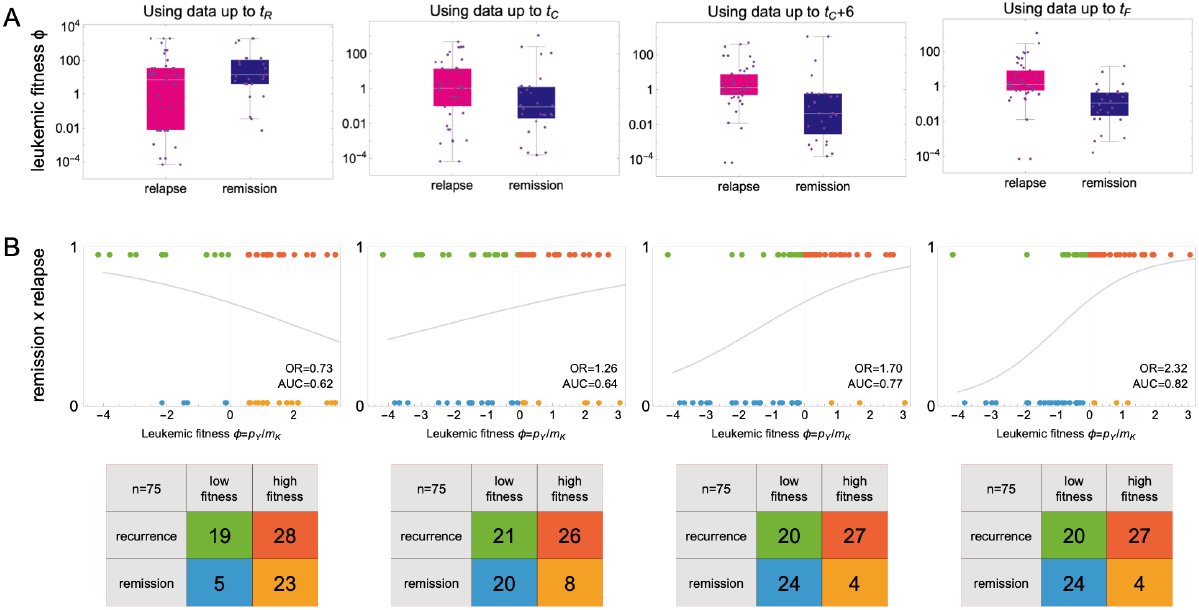
Leukemic fitness as a predictor of treatment outcome. **(A)** Distribution of estimated leukemic fitness *ϕ* = *p*_*Y*_/*m*_*K*_ (from the best-fit parameter) for patients grouped by simulated *reference outcome* (recurrence vs remission at 36 months after cessation using the best fit obtained with data up to *t*_*F*_), across four data selection strategies: leukemic fitness estimated with data up to dose reduction (*t*_*R*_), up to cessation (*t*_*C*_), up to six months after cessation (*t*_*C*_ + 6), and using the complete time course (*t*_*F*_). Fitness is shown on a log scale. The separation between groups increases progressively with data availability (p=0.173, 0.051, 1.4×10^−4^, and 9.6×10^−6^, respectively; Mann–Whitney U test), indicating that leukemic fitness becomes an increasingly discriminative predictor as more dynamic information is incorporated. **(B)** Logistic regression of treatment outcome (0 = remission, 1 = recurrence) as a function of estimated leukemic fitness *ϕ*, for each data selection strategy (*t*_*R*_, *t*_*C*_, *t*_*C*_+ 6, *t*_*F*_). Each point represents a patient; the fitted logistic curve is overlaid. The curve is decreasing for the shortest window; for the later windows it is increasing and becomes progressively steeper as the data range expands, reflecting the growing predictive power of fitness estimates. The odds ratio (OR, shown in each panel) per tenfold increase in fitness (logistic regression on log10 fitness), with 95% confidence interval, is 0.73 (0.55–0.97), 1.26 (0.96–1.65), 1.70 (1.20–2.40) and 2.32 (1.41–3.83) for the four windows, respectively. The area under the ROC curve derived from each logistic regression (AUC, shown in each panel) quantifies the ability of leukemic fitness to discriminate recurrence from remission outcomes as a continuous variable; values progress from 0.62 (data up to *t*_*R*_) to 0.64 (*t*_*C*_), 0.77 (*t*_*C*_ + 6), and 0.82 (*t*_*F*_), mirroring the progressive improvement in group separation observed in panel A; the value for the shortest window corresponds to an inverted association (OR below 1). Each panel further includes the 2×2 confusion matrix obtained with *ϕ* = 1 as the decision threshold, so that the four windows can be compared side by side: columns indicate the fitness-based prediction (remission: *ϕ* < 1; recurrence: *ϕ* > 1) and rows the simulated *reference outcome* at 36 months after cessation. Cell colors correspond to the patient point colors in the same panel, allowing individual patients to be identified across the two representations.

Casting the problem as a binary classification (fitness *ϕ* above vs below 1) and fitting a logistic regression to the fitness-outcome pairs (with fitness on a log10 scale) shows a progressive change in the association. The odds ratio (OR) per tenfold increase in fitness is 0.73 (95% confidence interval 0.55-0.97) with data up to *t*_*R*_, 1.26 (0.96-1.65) at *t_F_*, 1.70 (1.20-2.40) at *t*_*C*_ + 6, and 2.32 (1.41-3.83) at *t*_*F*_ (Fig 5B). The AUC takes the values 0.62, 0.64, 0.77 and 0.82 over the same four windows, following the same trend as the ensemble-based ROC analysis of Figs 2C and 3, although with lower values. At the shortest window, the logistic curve is decreasing (OR below 1), so the AUC of 0.62 reflects discrimination in the inverse direction, in line with the non-identifiability of the proliferation and immune kill rates under full-dose treatment alone (see Discussion). From the cessation window onwards the odds ratios are above 1 and increase with the amount of data. Together, these complementary perspectives confirm that leukemic fitness carries predictive information that increases substantially with the amount of data available, and that the transition from dose reduction to post-cessation dynamics provides the decisive signal.

A cross-tabulation of fitness-based prediction (Fig 5B, made with increasing estimation windows) against the *reference outcome* yields 27 true positives (high fitness, recurrence), 24 true negatives (low fitness, remission), 4 false positives (high fitness, remission), and 20 false negatives (low fitness, recurrence), corresponding to 51 correct predictions out of 75 patients (Fig 5B, fourth panel). This accuracy is inherently lower than the perfect discrimination reported above for the same data (Fig 2C), because the two criteria classify on diffierent grounds: the *reference outcome* scores simulated trajectories at a dated horizon (36 months post cessation), while the fitness-based thresholding at *ϕ* = 1 used here scores only the asymptotic attainability of cure and is blind to the position of the trajectory. The same cross-tabulation for the truncated windows is shown in the insets of the remaining panels: the two concordant cells hold 33 patients at *t*_*R*_, 46 at *t*_*C*_ and 51 at *t*_*C*_ + 6, so that concordance increases as the estimation window lengthens, reaching a plateau between the last two windows, in step with the group separation of panel A and the increasing odds ratios of the logistic regressions in panel B.

In each estimation window, it is the two discordant cells whose labels warrant careful interpretation, especially the predictions made with the full data (*t*_*F*_, rightmost panel in Fig 5B). A “false positive” (high fitness yet observed remission) does not represent a model error in the traditional sense: it identifies a patient whose leukemic fitness predicts eventual recurrence, but who has not yet been observed long enough for recurrence to manifest at the horizon of *t*_*C*_ + 36. This happens because the fitness prediction is asymptotic and states that recurrence will occur, not when. A patient with *ϕ* > 1 may therefore still be in apparent remission at *t*_*C*_ + 36 before tumor load eventually rises above the recurrence threshold (Fig 6A). Conversely, a “false negative” (low fitness yet observed recurrence) may reflect a patient who was treated for insufficient time and would have achieved durable remission under extended therapy (Fig 6B). Thus, rather than prediction errors, these discordant cases encode clinically meaningful information: the 20 patients with low fitness and observed recurrence require treatment extension until the separatrix is crossed (Fig 6B), while the 4 high-fitness patients with observed remission were likely not followed long enough for recurrence to manifest (Fig 6A). These two cells are clinically relevant, and we suggest respective interventions below, reframing the classification in terms of measurable quantities at the decision point rather than with respect to the *reference outcome* used here.

**Fig 6.**
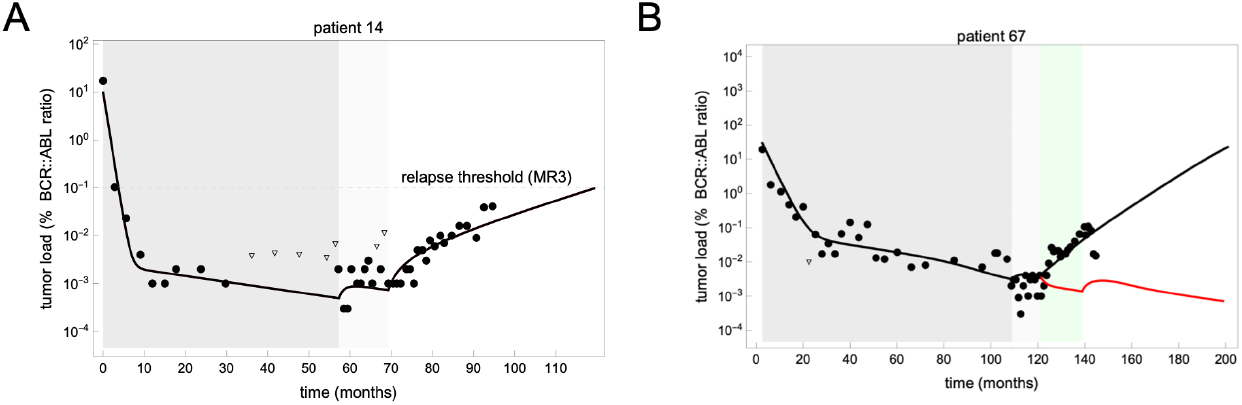
Treatment recommendations defined by estimated fitness and predicted outcome, illustrated in individual patients. Each panel shows one representative patient, together with the recommendation that follows from the group to which the estimated fitness and the predicted outcome assign that patient. **(A)**High estimated fitness with predicted remission: representative patient illustrating a discordant case with high fitness (*ϕ* > 1) but no observed recurrence at the 36-month post-cessation horizon. The best-fit trajectory (black) shows continued tumor growth, consistent with eventual recurrence. This case exemplifies why high-fitness patients classified as “false positives” in a short follow-up may represent true recurrences observable only at later time points. The recommendation for this group is therefore not to wait for the recurrence to be observed: since the estimated fitness already implies that it will occur, the patient is a candidate for the same maintenance strategy used for patients with high fitness and predicted recurrence, started in anticipation rather than in response. **(B)** Low estimated fitness with predicted recurrence: representative patient illustrating a discordant case with low fitness (*ϕ* < 1) but observed recurrence. The black trajectory shows the best-fit simulation under the standard protocol, in which treatment is discontinued at *t*_*C*_, resulting in molecular recurrence. The red trajectory shows the model-predicted outcome under an extended treatment protocol: resuming full-dose TKI at *t*_*C*_ for an additional 18 months before a second cessation attempt. In this scenario, the extended therapy is sufficient for the trajectory to cross the separatrix and enter the basin of attraction of *E*_0_, leading to sustained treatment-free remission. This illustrates how fitness-guided treatment extension could convert a patient who would otherwise experience recurrence into a TFR success, consistent with the separatrix crossing times estimated in Fig 7.

In summary, leukemic fitness *ϕ* integrates the balance between leukemic proliferative capacity and immune control into a single quantitative parameter. As a diagnostic criterion, it does not reproduce what was observed during follow-up; it reveals the system’s asymptotic fate. This retrospectively identifies which patients would benefit from extended treatment and which require ongoing maintenance therapy to control residual disease. We next investigate how fitness estimated from pre-cessation data can be used prospectively to guide these personalized treatment decisions.

### Fitness-guided treatment strategies: extending therapy toward remission and maintaining control in immune-insufficient patients

The preceding analysis established leukemic fitness *ϕ* as a single-parameter criterion for classifying patients into two distinct groups: those with *ϕ* < 1, for whom the cure equilibrium is stable and durable TFR is achievable, and those with *ϕ* > 1, for whom the immune system is insufficient to control residual disease and recurrence is the inevitable asymptotic outcome. Prospectively, however, the relevant classification combines two quantities that are both available at the decision point: the estimated fitness, below or above 1, and the outcome predicted at *t*_*C*_ + 36, remission or recurrence, yielding four patient groups, each implying diherent interventions:

i. For patients with low fitness and predicted remission, the trajectory has already crossed the separatrix and the remission is correctly predicted; no intervention is required.
ii. For patients with low fitness and predicted recurrence, durable remission is attainable but the separatrix has not yet been reached, so full-dose therapy should be extended until it is (Fig 6B).
iii. For patients with high fitness and predicted recurrence, recurrence cannot be averted by any finite treatment duration, and the relevant question becomes the maintenance dose that keeps residual disease under stable control.
iv. For patients with high fitness and predicted remission, the predicted remission is transient (Fig 6A), since the estimated fitness implies eventual recurrence, so the same maintenance strategy applies, initiated in anticipation rather than in response to an observed recurrence.

Since both coordinates are estimated from the same truncated data, a patient may be assigned to different groups depending on whether the estimate is made at *t*_*C*_ or at *t*_*C*_ + 6. From now on, we focus on these two estimation windows: the data up to *t*_*R*_ alone provide no meaningful information and considering treatment-adaptations after *t*_*C*_ + 6 appears too late.

For each of the two estimation windows (*t*_*C*_ and *t*_*C*_ + 6), we ask a paired question: given the estimated fitness *ϕ*, the predicted trajectory from the data available at that point, and the recommended intervention, what actually happens once that recommendation is simulated using the patient’s parameter set behind the *reference outcome* (defined above) subject to the modified treatment schedule? In other words, recommendation and evaluation are kept strictly separate: the group assignment, and hence the recommendation, uses only data up to *t*_*C*_ or *t*_*C*_ + 6, while the subsequent simulations use the best-fit parameters obtained from the complete patient time course (*t*_*F*_), essentially “the best knowledge” that we have about the patient. This design mimics the real clinical scenario: the clinician acts on the information available at the decision point, but the patient’s actual response unfolds according to their true underlying biology, so that any discrepancy between estimated and true fitness propagates into the simulated outcome.

For patients classified as low-fitness but predicted to experience recurrence because they did not reach the separatrix, the bifurcation analysis predicts that sustained remission is achievable through continued full-dose TKI therapy until the separatrix is crossed. Using data up to *t*_*C*_, 13 patients were classified into this category; using data up to *t*_*C*_ + 6, 22 patients were identified (here, the predicted outcome is obtained from the truncated data, not from the reference outcome). For each of these patients, we computed the additional full-dose treatment time required to cross the separatrix. The resulting distributions (Fig 7A) reveal substantial inter-patient heterogeneity, but also a systematic shift with the later decision point: the median additional treatment time is 31 months when treatment is resumed at *t*_*C*_, with 23%, 46%, and 54% of patients reaching the separatrix within one, two, and three years, respectively. When the decision is instead made at *t*_*C*_+ 6, the median additional treatment time drops to 12 months, with 55%, 59%, and 68% of patients reaching the separatrix within one, two, and three years respectively. The shift reflects a gain in accuracy rather than in outcome. With data only up to *t*_*C*_, the immune response has not yet been challenged by residual disease in the absence of TKI, so the fits are biased toward a scenario with little immune contribution, and therefore toward higher fitness and longer required treatment; the six additional months of post-cessation data pull the fitness estimate downward, bringing the predicted treatment time closer to the value implied by the patient′s own parameters. The advantage is that the patient is better characterized; and it would remain an advantage even if the corrected estimate were to turn out longer rather than shorter.

**Fig 7.**
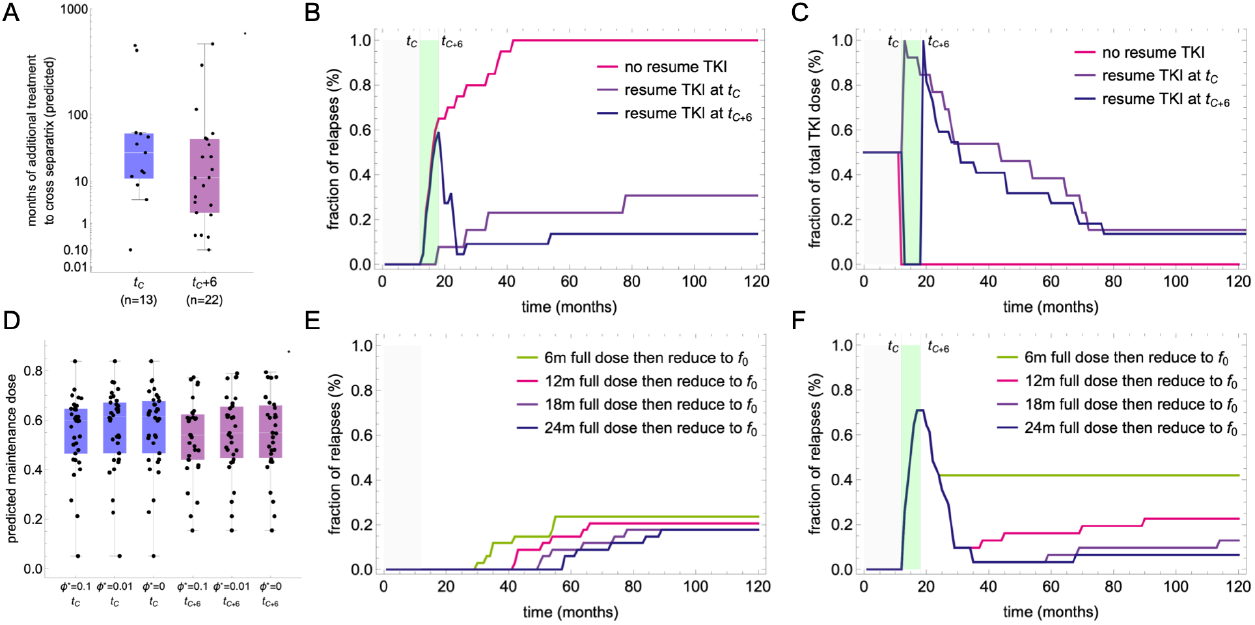
Fitness-guided adaptive treatment strategies for low- and high-fitness patients. **(A)** Distribution of the additional full-dose TKI treatment time predicted to be required for each low-fitness patient (*ϕ* < 1) to cross the separatrix, estimated using the best-fit parameters obtained from data up to *t*_*C*_ (n=13) or *t*_*C*_+ 6 (n=22). Each point represents one patient; boxes show the interquartile range with the median indicated by the horizontal line. **(B)** Cumulative fraction of patients predicted to experience recurrence over time under three scenarios: standard cessation with no treatment resumption (magenta), resumption of full-dose TKI at *t*_*C*_ for the individually predicted additional treatment time (purple), and resumption at *t*_*C*_+ 6 (dark blue). Shaded regions indicate the dose-reduction period before cessation at *t*_*C*_ (gray) and the six-month treatment-free window between *t*_*C*_ and *t*_*C*_+ 6 (green). All simulations use the best-fit parameters obtained from the complete patient time course (up to *t*_*F*_), so that the simulated outcome reflects the patient′s true underlying dynamics, while the decision to resume treatment, and its duration, is based on partial data (up to *t*_*C*_ or *t*_*C*_+ 6). **(C)** Fraction of total TKI dose administered over time, relative to a reference of continuous full-dose treatment, for the same three scenarios as in panel B. Despite the resumption of full-dose therapy, both adaptive strategies converge to a substantially reduced cumulative dose burden compared to continuous standard therapy, reflecting the transient nature of the additional treatment period required to cross the separatrix. **(D)** Distribution of the maintenance dose fraction *f*^*^ required for each high-fitness patient (*ϕ* > 1, estimated from data up to *t*_*C*_ or to *t*_*C*_+ 6) to achieve a target effective fitness *ϕ*^*^ ∈ {0.1, 0.01,0}. The dose fraction is computed analytically from the patient-specific parameter estimates as *f*^*^ = (*p*_*Y*_ – *m*_*K*_*ϕ*^*^)/(*q* + *p*_*Y*_). Each box shows the interquartile range across patients; individual points are overlaid. Despite substantial inter-patient variability in underlying parameters, the required maintenance doses cluster consistently between 50% and 60% of the standard dose across all target fitness values. **(E–F)** Cumulative fraction of high-fitness patients (*ϕ* > 1) predicted to experience recurrence over time under personalized maintenance therapy, for decision strategies *t*_*C*_ **(E)** and *t*_*C*_+ 6 **(F)**. In each scenario, full-dose TKI is resumed at the decision point for a re-induction period of 6, 12, 18, or 24 months (green, magenta, purple, and dark blue, respectively), after which the dose is reduced to the patient-specific maintenance dose *f*_0_. The green shaded region in panel F indicates the six-month treatment-free window preceding the *t*_*C*_+ 6 decision point. All simulations use the best-fit parameters from the complete patient time course (*t*_*F*_), so that the simulated outcomes reflect true patient dynamics while decisions are based on partial data. Re-induction periods of 12 months or longer suppress long-term recurrence to 10–25% under strategy *t*_*C*_, and to below 10% under strategy *t*_*C*_+ 6 with sufficient re-induction, demonstrating that personalized maintenance can achieve durable disease control in high-fitness patients.

The cohort-level impact of applying these individually predicted treatment extensions is shown in Fig 7B,C, where simulations use the best fit based on all available data (up to *t*_*F*_) for each patient, while the retreatment decision is made at *t*_*C*_ or *t*_*C*_+ 6 based on the data and the *ϕ* estimated until these time points. Compared to standard cessation without treatment resumption, under which all low-fitness patients eventually experience recurrence because the separatrix was not reached, resuming TKI for the individually estimated duration substantially reduces recurrence: 36 months after *t*_*C*_, 31% of patients experience recurrence under the *t*_*C*_ strategy and only 14% under the *t*_*C*_+ 6 strategy (Fig 7B). The recurrences that remain are expected: they occur in patients classified as low-fitness from partial data whose fitness, estimated from the complete time course used for the simulations, is in fact above 1, so that no finite extension of full-dose therapy can prevent recurrence. Despite this added treatment period, the cumulative dose burden remains modest: only 15% of patients are still receiving TKI five years after *t*_*C*_ under either strategy, and the fraction of total dose administered converges to similarly low levels for both approaches (Fig 7C). The *t*_*C*_+ 6 strategy, despite an initial six-month delay during which transient recurrences can occur, ultimately outperforms the *t*_*C*_ strategy on both recurrence rate and treatment burden, reflecting the fact that six additional months of data yield a better-informed decision. Together, these results suggest that a substantial proportion of patients currently classified as experiencing recurrence under standard cessation protocols could achieve durable remission through a personalized, fitness-guided extension of full-dose therapy.

For patients classified as high-fitness (*ϕ* > 1, independent of a predicted remission or recurrence) at *t*_*C*_ or *t*_*C*_+ 6, continued standard therapy maintains disease suppression but cannot lead to cure, as the cure equilibrium *E*_0_ is unstable regardless of treatment duration. For these patients, indefinite full-dose TKI therapy carries cumulative toxicity and quality-of-life costs without prospect of remission. The model offers a principled alternative: identifying the minimum maintenance dose sufficient to stabilize residual disease at a controlled level.

To formalize this, we introduce the concept of dose-dependent effective fitness. Assuming a linear dose-response relationship, a maintenance dose fraction *f* ∈ [0,1] relative to the standard dose yields an effective TKI kill rate of *fe*_*TKI*_, and the effective leukemic fitness under maintenance becomes

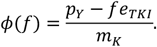

To achieve a certain target fitness *ϕ*^*^, we can calculate the required maintenance dose fraction *f*^*^. Solving *ϕ*(*f*^*^) = *ϕ*^*^ for a target fitness *ϕ*^*^ and substituting *e*_*TKI*_ = *q* + *p*_*Y*_ gives the required maintenance dose fraction

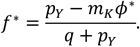

Since *q* is estimated in the first fitting step, and *p*_*Y*_ and *m*_*K*_ are estimated from data up to *t*_*C*_ or *t*_*C*_+ 6, this fraction *f*^*^ can be computed at the decision point to inform a personalized, model-based maintenance recommendation.

Using the best-fit parameter pairs estimated from data up to *t*_*C*_ and *t*_*C*_+ 6, we computed the maintenance dose fractions required to achieve target fitness values of *ϕ*^*^ = 0.1 and 0.01 across all high-fitness patients (Fig 7D). Despite inter-patient variability, the required dose fractions *f*^*^ increase and saturate as the target fitness *ϕ*^*^ decreases, with median values stabilizing around *f*^*^ ≈ 60% of the standard dose for strategy *t*_*C*_ and 55% for strategy *t*_*C*_+ 6. A natural limiting case arises when *ϕ*^*^ → 0, giving

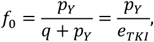

which corresponds to the dose fraction at which the TKI-induced kill rate exactly offsets leukemic proliferation in the absence of immune control, a biologically interpretable maximal required maintenance dose. The distributions of *f*_0_ across patients closely mirror those for *ϕ*^*^ = 0.01, with median values near 60% and 55% of standard dose for strategies *t*_*C*_ and *t*_*C*_+ 6, respectively. Given the small quantitative difference between *f*_0_ and the doses corresponding to lower target fitness values, and because *f*_0_ represents the conservative upper bound on required maintenance, we adopt *f*_0_ as the maintenance dose for all subsequent simulations.

Because high-fitness patients will have accumulated elevated tumor loads during the dose-reduction and treatment-free periods preceding *t*_*C*_ or *t*_*C*_+ 6, a period of full-dose re-induction is necessary before transitioning to the maintenance dose *f*_0_. We simulated re-induction periods of 6, 12, 18, and 24 months of full-dose TKI followed by long-term maintenance at *f*_*_, evaluating the resulting recurrence rates under both decision strategies (Fig 7E,F). For strategy *t*_*C*_, re-induction periods of 12 months or longer suppress recurrence to 10–25% over more than 5 years, with longer re-induction providing only marginal additional benefit (Fig 7E). For strategy *t*_*C*_+ 6, the initial six-month treatment-free window causes a transient spike in recurrences, but longer re-induction periods yield substantially better outcomes, with 18–24 months of re-induction reducing long-term recurrence to below 10% (Fig 7F).

Taken together, a fitness-guided adaptive protocol based on classifying patients at *t*_*C*_ or *t*_*C*_+ 6, extending therapy for low-fitness patients, and initiating personalized maintenance for high-fitness patients, is predicted to improve outcomes across the cohort while minimizing unnecessary treatment burden.

### Simulating maintenance therapy with population-level reduced doses in immune-insufficient patients

While for patients with a fitness *ϕ* > 1 an individually optimal maintenance dose *f*_*_ can be computed, the administration of this dose might be limited by practical constrains such as tablet size or frequency of intake. To assess the robustness of our findings in a clinical setting, we additionally simulated uniform population-level maintenance doses approximating clinically feasible regimens: 50% of standard dose (daily half-tablet), 57% (four doses per week), and 71% (five doses per week, mimicking treatment holidays on weekends). Under strategy *t*_*C*_, a maintenance dose of 50% leads to approximately 40% recurrence at 10 years regardless of re-induction duration, confirming that this dose is insufficient to stabilize disease in a substantial fraction of high-fitness patients (Fig 8A). This is consistent with our earlier report that half of the standard dose preserves molecular remission in about 87% of patients [16], which was obtained with the linear model without immune control (equation (1)) and applied to the cohort as a whole: the recurrence quoted here is confined to the high-fitness subgroup, precisely the patients in whom immune control is insufficient. Increasing the maintenance dose to 57% reduces recurrence to 25– 30%, with modest improvement for longer re-induction (Fig 8B). At 71%, recurrence falls to approximately 10% even with only 3 months of re-induction, approaching the performance of the personalized *f*_0_ protocol (Fig 8C). Under strategy *t*_*C*_+ 6, outcomes are generally worse for short re-induction periods due to the transient treatment-free window but improve substantially with longer re-induction: at 71% maintenance dose and 6-18 months of re-induction, long-term recurrence drops to approximately 15% (Fig 8D–F). Together, these results indicate that five doses per week represents a practical population-level maintenance regimen that approximates the efficacy of personalized dosing, while remaining feasible within standard clinical workflows.

**Fig 8.**
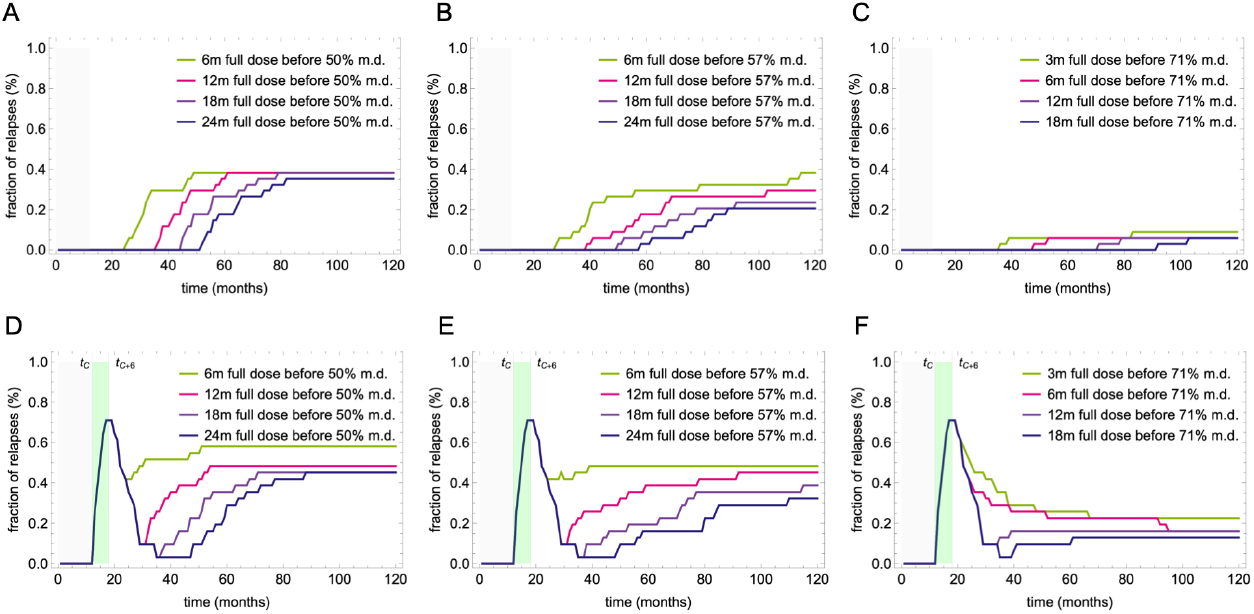
Recurrence rates under uniform population-level maintenance doses for high-fitness patients. Cumulative fraction of high-fitness patients (*ϕ* > 1) predicted to experience recurrence over time under uniform (non-personalized) maintenance doses (m.d.) of 50% **(A, D)**, 57% **(B, E)**, and 71% **(C, F)** of the standard TKI dose, for decision strategies *t*_*C*_ **(A–C)** and *t*_*C*_+ 6 **(D–F)**. In each panel, curves correspond to re-induction periods of 6, 12, 18, and 24 months of full-dose TKI (3, 6, 12 and 18 months for the 71% panels) prior to transitioning to the maintenance dose, as indicated in the legend. The green shaded region in panels D–F marks the six-month treatment-free window between the cessation time *t*_*C*_ and the decision point *t*_*C*_+ 6*m*. All simulations use the best-fit parameters from the complete patient time course. Maintenance at 50% of standard dose results in approximately 40% long-term recurrence regardless of re-induction duration under both strategies, indicating insufficient disease control for a substantial fraction of patients. At 57%, recurrence is reduced to 25–30% with longer re-induction. At 71% (five doses per week), long-term recurrence falls to approximately 10% under strategy *t*_*C*_, and to similarly low levels under strategy *t*_*C*_+ 6 with re-induction of 6 months or longer, approaching the performance of the personalized maintenance protocol (Fig 7E,F). These results suggest that five doses per week represents a practical population-level maintenance regimen that closely approximates the efficacy of individually computed doses.

These results confirm that, even for patients for whom cure is not achievable, a population-level maintenance protocol can provide durable disease control with substantially reduced treatment burden compared to indefinite full-dose therapy.

## Discussion

The central finding of this work is that the outcome of treatment-free remission is governed, in the model, by a single quantity that can be estimated from routine monitoring data: the leukemic fitness, defined as the ratio of the proliferative aggressiveness of residual leukemic cells to the anti-leukemic killing capacity, which can be estimated without explicitly representing the underlying immunology. The underlying biology is undeniably complex: no immune population or molecular marker has proven sufficient to predict TFR on its own, and remission maintenance likely reflects contributions from multiple compartments (NK cells, T cells, the bone marrow microenvironment, and cell-niche interactions) whose relative weights vary across patients. This complexity, however, need not be represented explicitly to capture the system-level behavior that matters for prediction. In fact, given the data these trials produce, it could not be resolved even if one tried: BCR::ABL1 time courses reflect the net effect of immune control on residual disease, but not its underlying composition. By encoding the collective anti-leukemic immune effect as a tumor load-dependent functional response, our suggested minimal model reproduces the qualitative patterns observed across patients while remaining analytically tractable. This suggests that, at the level of population dynamics relevant for TFR prediction, the net immune contribution behaves as a low-dimensional process: the critical unknown is not the identity of the immune effectors, but the balance between their collective killing capacity and the intrinsic proliferative rate of residual leukemic cells.

While complex models can incorporate multiple immune populations, pathways, and cytokine-mediated interactions, their predictive power is often hampered by parameter identifiability. The larger the model, the greater the likelihood of multiple parameter sets fitting the same data equally well yet corresponding to fundamentally different mechanistic explanations and divergent outcome predictions. Working through these increasingly sophisticated models exposes a persistent challenge: the inability to disentangle the intrinsic tumor proliferation rate (*p*_*Y*_) from the maximum immune kill rate (*m*_*K*_) using only data from patients under standard-dose TKI treatment. The antagonistic effects of these two processes are effectively masked by the potent action of the drug. Our minimal model does not resolve this non-identifiability, but it isolates it: by collapsing the immune component into a single effective kill rate, it reduces the question to the balance between the tumor’s aggressiveness (*p_Y_*) and the immune system’s control (*m_K_*), making the central obstacle for TFR prediction explicit rather than concealed within a high-dimensional parameter space. This reinforces the idea that predictive accuracy may require adaptive treatment protocols, designed to probe the system and disentangle the opposing forces that govern long-term tumor persistence versus remission.

The bifurcation analysis of the minimal model offers a mechanistic explanation for the well-established clinical observation that the duration of deep molecular response before cessation is among the strongest predictors of TFR success; specifically, total time on TKI and, to a lesser extent, time spent in MR4 [34,35]. In the model, this reflects the role of the quiescent LSC reservoir as the rate-limiting barrier to cure. Even when active tumor cells are suppressed below detectable levels, a quiescent pool persists and continuously reseeds the active compartment. The time required to deplete this reservoir below the separatrix threshold and thereby enter the basin of attraction of the cure equilibrium is determined jointly by the leukemic fitness *ϕ* and the quiescent cell activation rate *p*_*XY*_, and can span years even for patients with favorable immune profiles (Fig 4F). This provides a quantitative rationale for why premature treatment cessation, even in sustained and deep remission, frequently results in recurrence: the system has not yet crossed the separatrix, and the residual quiescent pool is sufficient to drive recurrence once TKI pressure is removed. Conversely, the model identifies a subset of patients (those with low fitness *ϕ* and high activation rate *p*_*XY*_) for whom the separatrix is reached relatively quickly, and who may be candidates for earlier, safer cessation attempts.

The implications of this finding are direct for the design of future clinical trials, monitoring strategies and adaptive treatment decisions. It suggests that the most valuable information for TFR prediction lies not in the steady-state response dynamics during treatment, but in the transient response to system perturbations [36]. Trials like DESTINY [31] or HALF [37], which include dose-reduction phases, are crucial because they generate precisely the kind of dynamic data needed to dissociate *p*_*Y*_ from *m*_*K*_. Our model provides a quantitative framework to assess how informative different perturbation strategies (e.g., different levels or durations of dose reduction) could be. The corollary is that better TFR prediction will come from better-designed data rather than from larger models.

An important methodological aspect of the present analysis is the strict separation between decision and evaluation: treatment strategies are derived from parameter estimates available at the time of clinical decision, while their outcomes are simulated using the best-fit parameters from the complete patient time course. This design propagates the uncertainty of the decision-time estimates into the predicted outcomes and thus provides a more realistic assessment than in-silico analyses in which the true parameters are assumed known. That the fitness-guided strategies perform well under this evaluation suggests that the approach is practically viable, a necessary condition for translation into clinical decision support.

Regarding fitness-guided treatment adaptations, our results suggest taking the treatment decision six months after cessation, when the post-cessation data allow a considerably more reliable estimate of leukemic fitness. Patients with low fitness whose treatment had been too short then would be assigned to resume full-dose TKI for an individually estimated, finite period until the separatrix is crossed. Patients with high fitness instead resume full-dose TKI for a re-induction period of about one year, and then continue on a reduced maintenance dose, either the patient-specific dose *f*_0_ (median 55–60% of the standard dose) or, as a practical uniform regimen, the standard dose on five days per week (71%).

Beyond probing the tumor-immune balance through dose alterations, the bifurcation analysis suggests an additional therapeutic option: pharmacologically increasing the quiescent LSC activation rate *p*_*XY*_ would accelerate depletion of the niche-bound reservoir and reduce the time required to cross the separatrix, potentially enabling earlier and safer treatment cessation. Interferon-alpha (IFN-α), which has been shown to mobilize quiescent HSCs and has been tested in combination with TKIs in CML [15,38,39], represents a candidate agent for this strategy, although the treatment schedules employed in existing trials were not specifically designed to maximize this effect, and the optimal timing and duration of such a combination remain to be determined. The model makes the idea testable: because the activation rate enters only through the speed at which the quiescent reservoir is emptied, a mobilizing agent should shorten the time needed to cross the separatrix without changing the leukemic fitness *ϕ* and should therefore be beneficial precisely for the low-fitness patients who are already destined for remission but have not yet reached it, while leaving high-fitness patients unaffected. Timing such a combination therapy in the months immediately preceding a cessation attempt, rather than spreading it across the whole treatment course, would be the most direct test of this prediction.

Several limitations should be acknowledged. First, the quiescent reservoir is treated as a single homogeneous compartment, whereas leukemic stem cells differ in depth of quiescence, niche dependence and drug sensitivity; a distribution of activation rates rather than a single value for *p*_*XY*_ would broaden the separatrix crossing times reported here and could reorder individual patients. However, as individual estimates are difficult to obtain, we retain the one-compartment description. Second, reducing the immune response to a function of tumor load absorbs every non-autonomous influence on immune efficacy (inhibitory signaling, effector exhaustion, niche-mediated protection, age and comorbidity, concomitant medication) into constants that do not vary in time, although immune competence plausibly drifts over the years spanned by these trajectories; such effects can be incorporated if patient-specific data become available. Third, parameters were estimated by Monte Carlo sampling of the parameter space, retaining point best fits; hierarchical or Bayesian inference with population-level priors would quantify uncertainty more faithfully and, by borrowing strength across patients, might constrain *p*_*Y*_ and *m*_*K*_ further than patient-by-patient fitting allows. We nevertheless retained the simple sampling procedure, identical to that of [18], because it allows a direct comparison with the full model and preserves a broad coverage of the parameter space. Fourth, the *reference outcome* against which classification performance is measured is itself model-derived and extrapolated beyond the observed follow-up, so the reported accuracies are conditional on the minimal model describing the untested regime adequately. A direct comparison with observed time to recurrence, accounting for censoring, was not feasible at a common horizon because postcessation follow-up is short and heterogeneous, and validation in cohorts with longer follow-up is required. Finally, the present approach is complementary to CML models that represent immune populations explicitly or that treat stem-cell dynamics stochastically [13,17,20]: the minimal model is a simpler description of the biology, but matched to what the available data can identify. Our approach is a pragmatic compromise: it trades mechanistic detail for identifiability and does not attempt to describe the full biology of CML remission, only to estimate the one quantity from routinely available BCR::ABL1 time courses that can actually predict outcome.

In conclusion, this work consolidates our earlier CML models into a single framework: it shows that the central predictive challenge for TFR reduces to estimating the balance between two key parameters, *p*_*Y*_ and *m*_*K*_, and that this balance, once estimated, directly informs both prognosis and treatment design. The leukemic fitness *ϕ* = *p*_*Y*_/*m*_*K*_ links the mathematical structure of the model to both patient heterogeneity and treatment decisions. The same strategy may be useful beyond CML. In many cancers, the processes that shape treatment response, such as immune surveillance or interactions with the microenvironment, cannot be measured directly. Condensing them into a few effective parameters that can be estimated from routinely collected data, as done here, may inspire other minimalist models in mathematical oncology: complex, unmeasurable phenomena can be expressed in simpler terms that still describe what is observed. Future work should focus on prospective validation of fitness-guided protocols, on developing monitoring strategies that can estimate *ϕ* earlier and more reliably, and on designing clinical trials that explicitly probe the tumor-immune balance through adaptive dose perturbations, toward predictive and personalized treatment of CML.

## Acknowledgments

ACF was supported by Alexander von Humboldt Foundation and CNPq.

## Author contributions

Conceptualization: ACF, IG.

Data curation: ACF.

Formal analysis: ACF, NWS.

Investigation: ACF.

Methodology: ACF.

Software: ACF.

Supervision: ACF.

Validation: ACF.

Visualization: ACF.

Writing – original draft: ACF.

Writing – review and editing: ACF, IG and NWS.

## Data availability statement

The patient data used in this study were previously published in references [18,27]. In this submission, these data and all author-generated code supporting the findings of this study are made available to reviewers in Zenodo via the link: https://zenodo.org/records/23163761?preview=1&token=eyJhbGciOiJIUzUxMiJ9.eyJpZCI6ImQzM2M4MTIyLWQ4ZDgtNGRkNi05ZmNkLTUwMTdlMTEwODI1NSIsImRhdGEiOnt9LCJyYW5kb20iOiIwMTI3NjA1MDBiODM2MDRlOGE4ZDRkOTA3ZmFlYmNiZCJ9.pYejh_3R5gzdDbVLwzpSzPrNzX-4Htf308GNl93psvQjE1bzFA4uR_v8eL1p_TbAvpmQzXutaPejFNcKOo_TEw

Upon acceptance, the data and the code will be made publicly available.

## Supporting information

***S1 Appendix. Mathematical analysis and supplementary figures***. Contains the complete stability and bifurcation analysis of the minimal model, together with Figs A–C.

