## Supplementary Material for "Leukemic fitness predicts treatment-free remission and guides adaptive therapy in chronic myeloid leukemia"

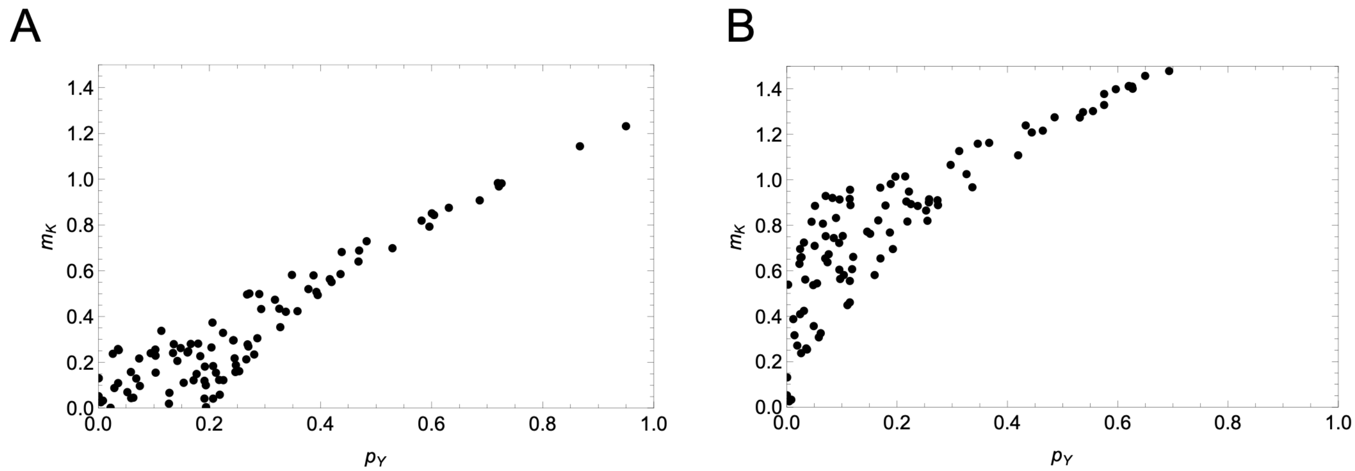


**Fig A. Compensatory ridge between proliferation and immune kill rate within the retained fit ensembles.** Scatter plots where each point is one of the 100 best-fitting parameter pairs for two representative patients (patient 14 in A, patient 15 in B), plotted as maximum immune kill rate $m_K$ against intrinsic proliferation rate $p_Y$. Fits align along a ridge along which an approximately proportional increase in proliferation is offset by a proportional increase in immune killing, so the data constrain the ratio direction more tightly than either rate individually. Across the 75 DESTINY patients the within-patient Spearman correlation has median ρ = 0.84 (IQR 0.66–0.93), is positive in every patient and significant at the 5% level in 74 of 75; the median total-least-squares slope is 1.28.

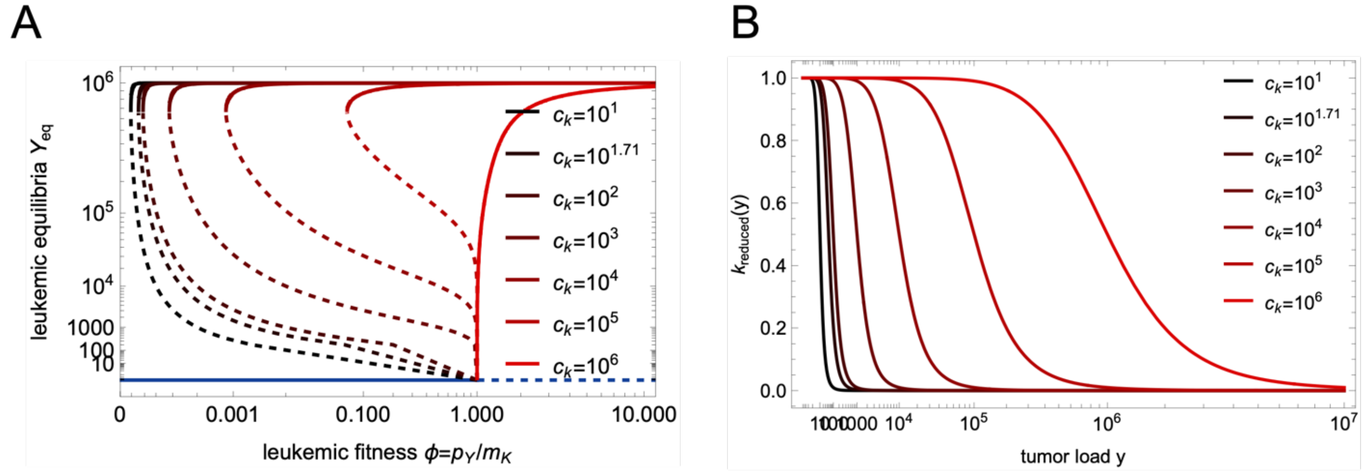


**Fig B. Bifurcation structure and immune kill rate for varying $c_K$. (A)** Bifurcation diagram showing the curve of leukemic equilibria $Y_{\text{eq}}$ as a function of leukemic fitness $\phi = p_Y/m_K$, for seven values of $c_K \in \{10^1, 10^{1.71}, 10^2, 10^3, 10^4, 10^5, 10^6\}$ (black to red). For each value of $c_K$, the diagram displays the stable branches (solid) and the saddle branch $E_L$ (dashed). The saddle-node bifurcation threshold $\phi_2$ increases with increasing $c_K$. The narrow regime $1 < \phi < \phi_1$ in which a third root $Y_D$ exists is not visible in the diagram, as $\phi_1 \approx 1 + 6.6 \times 10^{-10}$ for the estimated $c_K = 10^{1.71}$, and the corresponding equilibrium $E_D$ lies below $Y = 1.3 \times 10^{-3}$, well beneath the resolution of the plot and the biological threshold of a single cell. **(B)** Per-capita immune kill rate $k_{\text{reduced}}(Y)$ as a function of tumor load $Y$, for the same values of $c_K$. Larger $c_K$ shifts the transition from immune suppression to immune activity toward higher tumor loads, corresponding to a scenario in which immune efficacy is restored at higher residual disease levels. The estimated value $c_K = 10^{1.71}$ (dark brown) places the immune activation threshold well below MR3 ($10^3$ cells), consistent with clinical observations of immune recovery during deep molecular response.

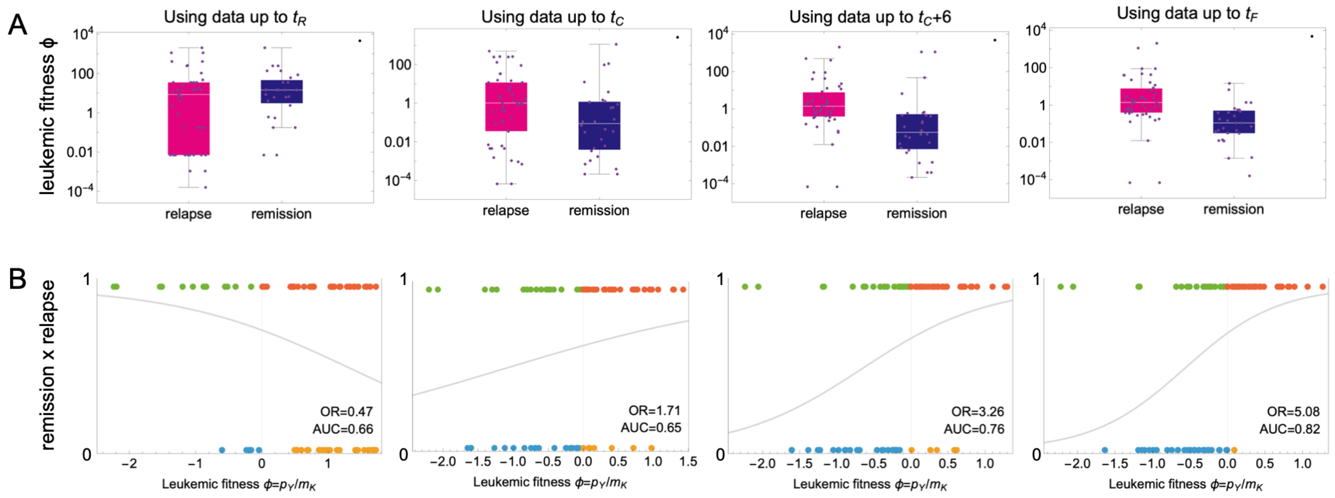


**Fig C. Leukemic fitness as a predictor of treatment outcome. (A)** Distribution of estimated *mean* leukemic fitness $\phi = p_Y/m_K$ (mean across all 100 best-fits) for patients grouped by simulated outcome (recurrence vs remission at 36 months after cessation), across four data selection strategies: up to dose reduction ($t_R$), up to cessation ($t_C$), up to six months after cessation ($t_C + 6$), and using the complete time course ($t_F$). Fitness is shown on a log scale. The separation between groups increases progressively with data availability (p=0.109, 0.049, $5.7\times10^{-4}$, and $4.8\times10^{-6}$, respectively; Mann–Whitney U test). **(B)** Logistic regression of treatment outcome (0 = remission, 1 = recurrence) as a function of estimated leukemic fitness $\phi$, for each data selection strategy ($t_R$, $t_C$, $t_C + 6$, $t_F$). Each point represents a patient; the fitted logistic curve is overlaid. The curve is decreasing for the shortest window; for the later windows it is increasing and becomes progressively steeper as the data range expands, reflecting the growing predictive power of fitness estimates. The odds ratio (OR, shown in each panel) per tenfold increase in fitness (logistic regression on log10 fitness), with 95% confidence interval, is 0.47 (0.25-0.88), 1.71 (0.92-3.15), 3.26 (1.50-7.11) and 5.08 (1.92-13.45) for the four windows, respectively. The area under the ROC curve derived from each logistic regression (AUC, shown in each panel) quantifies the ability of leukemic fitness to discriminate recurrence from remission outcomes as a continuous variable; values are 0.66 (data up to $t_R$) to 0.65 ($t_C$), 0.76 ($t_C + 6$), and 0.82 ($t_F$), mirroring the progressive improvement in group separation observed in panel A; the value for the shortest window corresponds to an inverted association (OR below 1).

### Mathematical Analysis

**Equilibrium points.** Because we are interested in the response after TKI stop, we analyze the long-term dynamics of the treatment-free system ($e_{TKI} = 0$): in this view, TKI acts as a transient external force that moves trajectories across the phase plane, but the asymptotic fate of the system (remission or recurrence) is determined entirely by the permanent dynamics and the position of the separatrix.

The non-trivial equilibria of the model are obtained by setting $\dot{Y} = 0$ (with $\dot{X} = 0$ implying $X = (p_{YX}/p_{XY})Y$), which yields the condition

$$\phi Y \left(1 - \frac{Y}{T_Y}\right) = k_{\text{reduced}}(Y)Y$$

where $\phi = p_Y/m_K$ is the leukemic fitness. One solution is always $Y = 0$, corresponding to the cure equilibrium $E_0$. The remaining equilibria satisfy

$$\phi\left(1-\frac{Y}{T_Y}\right)\left(1+\frac{Y^2}{c_K^2}\right)=1, \tag{A1}$$

or equivalently, the cubic polynomial

$$a_0+a_1Y+a_2Y^2+a_3Y^3=0$$

with coefficients

$$a_0=1-\phi,\ a_1=\frac{\phi}{T_Y},\ a_2=-\frac{\phi}{c_K^2},\ a_3=\frac{\phi}{c_K^2T_Y}.$$

**Counting positive roots via Descartes' rule**. We have two cases for the sign of $\phi-1$ :

- Low fitness ($0<\phi<1$): the sign sequence of $(a_0,a_1,a_2,a_3)$ is (+,+,-,+), yielding 0 or 2 positive roots, denoted $Y_H>Y_L>0$ when they exist. Substituting $Y\rightarrow -Y$ gives sign sequence (+,-,-,-), yielding exactly 1 negative root (biologically irrelevant).
- High fitness ($\phi>1$) : the sign sequence is (-,+,-,+), yielding 1 or 3 positive roots. When a single positive root exists, it is denoted $Y_H>0$; when three exist, they are denoted $Y_H>Y_L>Y_D>0$. Substituting $Y\rightarrow -Y$ gives sign sequence (-,-,-,-), yielding no negative roots.

**Bifurcation curve**. To analyze the existence and number of positive roots in detail, we solve equation (A1) for $\phi$ and express $\phi$ as a function of $Y$ along the equilibrium curve

$$\phi_{\text{eq}}(Y)=\frac{c_K^2T_Y}{(T_Y-Y)(c_K^2+Y^2)}$$

which is positive if and only if $0\leq Y<T_Y$, with a vertical asymptote at $Y=T_Y$ close to the high-load root $Y_H$. The bifurcation diagram obtained by plotting $Y$ as a function of $\phi$ for several values of $c_K$ is shown in Fig. B.

**Saddle-node bifurcations.** The local extrema of $\phi_{\text{eq}}(Y)$, where $\phi'_{\text{eq}}(Y)=0$, correspond to saddle-node bifurcations. Computing the derivative yields

$$\phi'_{\text{eq}}(Y)=\frac{c_K^2T_Y(c_K^2-2T_YY+3Y^2)}{(T_Y-Y)^2(c_K^2+Y^2)^2}.$$

The zeros of $\phi'_{\text{eq}}(Y)$ are the roots of the quadratic $3Y^2-2T_YY+c_K^2=0$, which has real solutions if and only if its discriminant is non-negative, i.e.,

$$c_K<\frac{T_Y}{\sqrt{3}}.$$

This condition is biologically reasonable: its violation would imply that the immune response threshold $c_K$ is comparable to the leukemic carrying capacity $T_Y$, corresponding to a scenario where immune activation is maximally effective only near full tumor burden - a situation inconsistent with the estimated parameter values (see Fig. B, which shows $k_{\text{reduced}}(Y)$ for several values of $c_K$ ). We therefore restrict our analysis to $c_K<T_Y/\sqrt{3}$, a biologically well-founded assumption under which $\phi'_{\text{eq}}(Y)$ has exactly two zeros, given by

$$y_1=\frac{T_Y-\sqrt{T_Y^2-3c_K^2}}{3},\ y_2=\frac{T_Y+\sqrt{T_Y^2-3c_K^2}}{3}$$

Writing $c_K = \varepsilon T_Y$ with $\varepsilon \ll 1$ (for the estimated value $c_K = 10^{1.71}$, we have $\varepsilon = 10^{1.71-6} \approx 5.1 \times 10^{-5}$ ), a series expansion to leading order gives

$$y_1 \approx \frac{\varepsilon^2 T_Y}{2} + O(\varepsilon^4),\ y_2 \approx \frac{2T_Y}{3} - \frac{\varepsilon^2 T_Y}{2} + O(\varepsilon^4).$$

The corresponding saddle-node bifurcation values of $\phi$ are $\phi_1 = \phi_{\text{eq}}(y_1)$ and $\phi_2 = \phi_{\text{eq}}(y_2)$, which expand to leading order as

$$\phi_1 \approx 1 + \frac{\varepsilon^2}{4} + O(\varepsilon^4),\ \phi_2 \approx \frac{27\varepsilon^2}{4} + O(\varepsilon^4),$$

with $\varepsilon = c_K/T_Y$.

**Complete bifurcation portrait**. The following regimes are identified as $\phi$ increases:

- $0 < \phi < \phi_2$: no non-trivial positive equilibria exist. The cure equilibrium $E_0$ is the unique attractor and is globally stable. For the estimated value $c_K = 10^{1.71}, \phi_2 \approx 27(c_K/T_Y)^2/4 \approx 1.78 \times 10^{-8}$.
- $\phi = \phi_2$: a saddle-node bifurcation occurs, creating two non-trivial equilibria $Y_H > Y_L > 0$. The high-load equilibrium $E_H$ is stable and the low-load equilibrium $E_L$ is a saddle (see stability analysis below). Bistability between $E_0$ and $E_H$ begins.
- $\phi_2 < \phi < 1$: bistability between $E_0$ and $E_H$, with the stable manifold of $E_L$ acting as the separatrix between their basins of attraction. This is the clinically relevant regime for treatment-free remission.
- $\phi = 1$: a transcritical bifurcation occurs. A third root $Y_D < Y_L$ emerges from $Y = 0$, giving rise to a stable deep-remission equilibrium $E_D$ at a very low tumor load. Simultaneously, $E_0$ loses stability and becomes a saddle. Mathematical cure ($Y = 0$) is no longer an attractor.
- $1 < \phi < \phi_1$: three positive roots $Y_H > Y_L > Y_D$ coexist. Bistability arises between $E_H$ and $E_D$, with $E_L$ separating their basins of attraction. For the estimated $c_K = 10^{1.71}, \phi_1 \approx 1 + (c_K/T_Y)^2/4 \approx 1 + 6.58 \times 10^{-10}$, and $Y_D$ at $\phi = 1$ takes the value $Y_L \approx 3.65 \times 10^{-3}$ - well below one cell. This regime is therefore mathematically distinct from $\phi > 1$ but biologically indistinguishable from it: complete cure is impossible, and $E_D$ represents a state of negligibly low residual disease that is not physiologically meaningful.
- $\phi = \phi_1$: a second saddle-node bifurcation, at which $Y_L$ and $Y_D$ collide (at $Y = y_1 \approx 1.32 \times 10^{-3}$ for $c_K = 10^{1.71}$ ) and annihilate each other.
- $\phi > \phi_1$: only $Y_H$ remains. $E_H$ is the unique attractor and is globally stable. Recurrence is the inevitable long-term outcome.

In summary, the biologically relevant bifurcation structure consists of a saddle-node bifurcation at $\phi_2 \ll 1$ (onset of bistability) and a transcritical bifurcation at $\phi = 1$ (loss of curability), with a narrow and biologically negligible intermediate regime between $\phi = 1$ and $\phi_1 \approx 1$. For all practical purposes, $\phi = 1$ constitutes the sharp threshold separating curable from incurable disease dynamics.

**Stability Analysis of Equilibria and Boundedness of Trajectories**

To analyze the stability of the equilibria, we rewrite the two-dimensional system as

$$\dot{X} = -p_{XY}X + p_{YX}Y,\ \dot{Y} = p_{XY}X - p_{YX}Y + Yu(Y),$$

where

$$u(Y) = p_Y\left(1 - \frac{Y}{T_Y}\right) - m_K k_{\text{reduced}}(Y)$$

Note that $u(Y) = 0$ if and only if $Y$ is a root of the equilibrium equation (A1). The Jacobian of the system is

$$J = \begin{pmatrix} -p_{XY} & p_{YX} \\ p_{XY} & -p_{YX} + u(Y) + Yu'(Y) \end{pmatrix}.$$

**Stability of $E_0$.** At the cure equilibrium $E_0 = (0,0)$, the Jacobian reduces to

$$J_0 = \begin{pmatrix} -p_{XY} & p_{YX} \\ p_{XY} & -p_{YX} + u(0) \end{pmatrix},$$

whose eigenvalues are the roots of $\lambda^2 - \text{tr}(J_0)\lambda + \det(J_0) = 0$. Since

$$\det(J_0) = p_{XY}(m_K - p_Y) = p_{XY}m_K(1 - \phi),$$

the following cases arise:

- If $\phi > 1$ : $\det(J_0) < 0$, so the eigenvalues are real with opposite signs. Therefore $E_0$ is a saddle point, and the cure equilibrium is unstable.
- If $\phi < 1$ : $\det(J_0) > 0$. Moreover, $\text{tr}(J_0) = -p_{XY} - p_{YX} + u(0) = p_Y - m_K - p_{XY} - p_{YX} < 0$, since $\phi < 1$ implies $p_Y < m_K$. Thus both eigenvalues have negative real part, and $E_0$ is locally asymptotically stable.

**Stability of non-trivial equilibria**. At a non-trivial equilibrium $E = (X_E, Y_E)$, with $X_E = (p_{YX}/p_{XY})Y_E$ and $u(Y_E) = 0$, the Jacobian reduces to

$$J_E = \begin{pmatrix} -p_{XY} & p_{YX} \\ p_{XY} & -p_{YX} + Y_E u'(Y_E) \end{pmatrix},$$

with

$$\det(J_E) = -p_{XY}Y_E u'(Y_E),\ \text{tr}(J_E) = -p_{XY} - p_{YX} + Y_E u'(Y_E).$$

The sign of $u'(Y_E)$ determines the local behavior:

- If $u'(Y_E) > 0$: $\det(J_E) < 0$, so the eigenvalues are real with opposite signs and $E$ is a saddle point.
- If $u'(Y_E) < 0$: $\det(J_E) > 0$ and $\text{tr}(J_E) = -p_{XY} - p_{YX} + Y_E u'(Y_E) < 0$, so both eigenvalues have negative real part and $E$ is locally asymptotically stable.

To determine the sign of $u'$ at each root, we note that $u(Y) < 0$ for sufficiently large $Y$ (since the logistic term $p_Y(1 - Y/T_Y)$ dominates and becomes negative, while $m_K k_{\text{reduced}}(Y) \geq 0$). Therefore, at the largest root $Y_H$, the function $u$ must be decreasing, i.e., $u'(Y_H) < 0$. It follows that $E_H$ is locally asymptotically stable whenever it exists. At the immediately preceding root $Y_L$, the function $u$ must be increasing, i.e., $u'(Y_L) > 0$, so $E_L$ is a saddle point whenever it exists. By the same argument, $E_D$ is locally asymptotically stable whenever it exists.

**Boundedness of trajectories**. We now show that all trajectories initiating in the non-negative quadrant $\{X \geq 0, Y \geq 0\}$ remain bounded. Define $\Sigma = 2X + Y$. Differentiating along solutions of the system and adding $\gamma\Sigma$, where $\gamma > 0$ will be fixed later, gives

$$\dot{\Sigma} + \gamma\Sigma \leq X(2 - p_{XY}\gamma) + AY - BY^2$$

where

$$A = 1 + p_Y\gamma + p_{YX}\gamma,\ B = \frac{p_Y\gamma}{T_Y}.$$

Since $AY - BY^2 \leq \delta := A^2/(4B)$ for all $Y \geq 0$, choosing $\gamma > 2/p_{XY}$ ensures that the coefficient of $X$ is negative, yielding

$$\dot{\Sigma} + \gamma\Sigma \leq \delta.$$

By Gronwall's inequality, this implies $\Sigma(t) \leq \Sigma(0)e^{-\gamma t} + \delta(1 - e^{-\gamma t}) \leq \max\{\Sigma(0), \delta\}$ for all $t \geq 0$. Since the nonnegative quadrant is forward-invariant under the flow of the system (as $\dot{X}\big|_{X=0} = p_{YX}Y \geq 0$ and $Y|_{Y=0} = p_{XY}X \geq 0$ ), it follows that both $X(t)$ and $Y(t)$ remain bounded for all $t \geq 0$.

**Exclusion of periodic orbits**. To prove that the model admits no periodic orbits in the non-negative quadrant, we use the theory of cooperative dynamical systems [Hirsch1982]. For planar cooperative irreducible systems, a classical result of Hirsch [Hirsch1982] states that the $\omega$-limit set of any precompact orbit consists of a single equilibrium; in particular, no non-constant periodic orbit can exist. To show that the model is cooperative and irreducible, note that the Jacobian of system (6) is

$$J = \begin{pmatrix} -p_{XY} & p_{YX} \\ p_{XY} & -p_{YX} + u(Y) + Yu'(Y) \end{pmatrix}.$$

The off-diagonal entries $J_{12} = p_{YX} > 0$ and $J_{21} = p_{XY} > 0$ are strictly positive constants, independent of the state variables. Therefore, the system is cooperative in the open positive quadrant $\{X > 0, Y > 0\}$ because all off-diagonal entries of $J$ are non-negative, and irreducible because the associated dependency graph is strongly connected (each variable directly influences the other). Since all trajectories are bounded (as shown above), every orbit is precompact. Therefore, every solution of the model converges to one of the equilibria identified in the bifurcation analysis above, and the long-term dynamics are entirely governed by their respective basins of attraction.
